# Guild-level response of the gut microbiota to isocaloric time-restricted feeding in high-fat diet-fed mice

**DOI:** 10.1101/2023.05.23.541832

**Authors:** Shreya Ghosh, Yue Li, Xin Yang, Guojun Wu, Chenhong Zhang, Liping Zhao

## Abstract

Time-restricted feeding (TRF) during the active phase protects against high-fat diet (HFD)-induced obesity, and its impact on gut microbiota has been previously investigated using bacterial taxa as functional units. However, in the gut ecosystem, bacteria from different taxonomic backgrounds form coherent functional groups called guilds, whose members exhibit co-abundant behavior. Thirty-five co-abundance groups (CAGs), clustered from 297 prevalent amplicon sequence variants (ASVs), showed greater concordance with beta-diversity plots based on all 1131 ASVs than the 130 classifiable genera, leading to a significantly improved preservation of community-level information. TRF-enriched CAGs positively correlated with metabolic improvement, while TRF-reduced CAGs negatively correlated. TRF restored the diurnal rhythm of most of these key CAGs. Novel ASVs, unclassifiable at the genus level, were identified in these key CAGs. Overall, this suggests that the key bacterial guilds may mediate the beneficial metabolic effects of TRF through the restoration of diurnal oscillation.

## Introduction

Ad libitum feeding of high-fat diet (HFD) to mice can induce obesity and related comorbidities, with disrupted gut microbiota playing a pivotal role in mediating the effects of the diet on body weight regulation and metabolism^1–3^. Mice are nocturnal and consume most of their food during the night (i.e., the active phase^4,5^). However, ad libitum access to HFD tends to alter the eating pattern in mice by increasing food intake during the daytime (i.e., rest phase^4,5^). Interestingly, the regime of time-restricted feeding (TRF), where access to food is limited to 8-12 hours during the active phase, alleviates HFD-induced weight gain and metabolic dysfunction in mice^5,6^. Despite many efforts, the underlying mechanism involved in TRF-induced improvement of the overall metabolic phenotype observed in HFD-fed mice is not completely understood.

Recent studies showed that the gut microbiota undergoes diurnal oscillation, and these oscillations are driven mostly by diet and feeding patterns in mice^7–9^. In HFD-induced obese mice, the diurnal oscillating nature of the gut microbiota is diminished compared to the mice fed a control diet^8^. Recent studies have shown that TRF modulates the gut microbiota in mice^8,10,11^. Mice fed a normal chow diet exhibited diurnal oscillations in their gut microbiota at the phylum level, with increased abundance of Firmicutes during the active phase, which decreases during the resting phase^8^. Similarly, Bacteroidetes and Verrucomicrobia levels increased during the resting phase and decreased during the active phase. In mice with ad libitum access to HFD, the diurnal oscillations at the phylum level were dampened, with the Firmicutes phylum being dominant at all timepoints compared to mice fed with a normal chow diet. TRF did not restore the diurnal oscillations at the phylum level. However, TRF did restore diurnal oscillations in some members of the gut microbiota. For example, TRF restored the diurnal oscillations in the *Lactobacillus* genus, which was lost in mice with ad libitum access to HFD. Another study reported that TRF increased Bacteroidetes abundance and decreased Firmicutes abundance^11^. Additionally, a study found that *Ruminococcus* and Christensenellaceae were enriched in the TRF group^10^. These studies all used various taxa from genus to phylum as units of microbial features during analysis. In such taxon-based analysis, the high-resolution reads obtained by sequencing a portion of the hypervariable region of the 16S rRNA gene are collapsed into a taxonomic. Novel sequences that cannot be assigned a desired level of taxon such as genus are discarded with no further analysis, leaving a partial and distorted picture of the microbiome. Additionally, taxon-level analysis assumes that members belonging to the same taxa have the same ecological response to environments. However, significant genomic diversity exists among bacterial strains that belong to the same named species, which can result in significant differences in their ecological functions^12,13^. Members from the same taxon may show opposite response to the same dietary intervention^14^. Some members in the same taxon may not respond to dietary intervention. Thus, combining ecologically competing bacteria into the same taxon or summing up responding strains with non-responding strains as one unit will lead to spurious results. Recently, a guild-based strategy has been adopted as an ecologically meaningful way to resolve the issues of high-dimensionality and sparsity of the microbiome data^12,15,16^. The gut microbiota is a complex ecosystem with its members interacting with one another and forming coherent functional groups termed “guilds”^17^. Members in the same guild cooperate with each to thrive or decline together, showing co-abundance behavior. Potential guilds can be identified by clustering prevalent ASVs based on their co-abundance patterns^15,18^. Guild-level abundance can be correlated with host bio-clinical parameters to identify functional groups in responding to dietary intervention and modulating host health-relevant phenotypes. In comparison to taxon-based analysis, the guild-based approach offers an ecologically sound tool to identify the key members of the gut microbial community associated with a particular host phenotype.

Moreover, the beneficial effects of a TRF regime were proportional to the duration of fasting, but often increasing the fasting duration results in a lower caloric intake^19–21^. So, some of the TRF-associated metabolic benefits observed so far may be mediated by reduced caloric intake. Studies showed that reduced calorie intake significantly modulated the gut microbiota^22^. Thus, a lower calorie intake may confound the impact of TRF on the gut microbiota. Taken together, our understanding is still limited on how the gut microbiota responds to TRF and contributes towards improved metabolic health in mice.

In this study, we examine the impact of TRF on the gut microbiota, glucose tolerance, and diurnal oscillation of the gut microbiota in mice fed a high-fat diet (HFD) with isocaloric intake with a normal fat diet as controls. Our aim was to determine if TRF could restore the diurnal oscillation of gut microbiota in HFD-fed mice and whether this restoration was associated with improved metabolic health. We clustered 297 prevalent amplicon sequence variants (ASVs) into 35 co-abundance groups (CAGs). We found that the beta-diversity PCoA plot based on the 35 CAGs better preserved the community level differences than that based on 130 genera. This indicates that guild-based analysis keeps most of the microbiome information intact. We then analyzed guild-level response of the gut microbiome to TRF. We identified key bacterial guilds which may mediate the effects of TRF on host health.

## Results

### TRF resulted in reduced body weight gain and glucose intolerance compared to ad libitum HFD feeding despite isocaloric intake

To assess if TRF influenced the observed metabolic outcomes independently of calorie intake, 8-week-old male C57BL/6J mice were randomly assigned to one of the following four groups (Fig. 1A): (i) the NA group was given ad libitum access to NFD, (ii) the NR group had time-restricted access to NFD, meaning access to food was limited to the active phase (ZT12-ZT0), (iii) the FA group was given ad libitum access to HFD, and (iv) the FR group had time-restricted access to HFD. Calorie intake remained the same for both treatments on the same diet (Fig. 1B).

**FIG 1.**
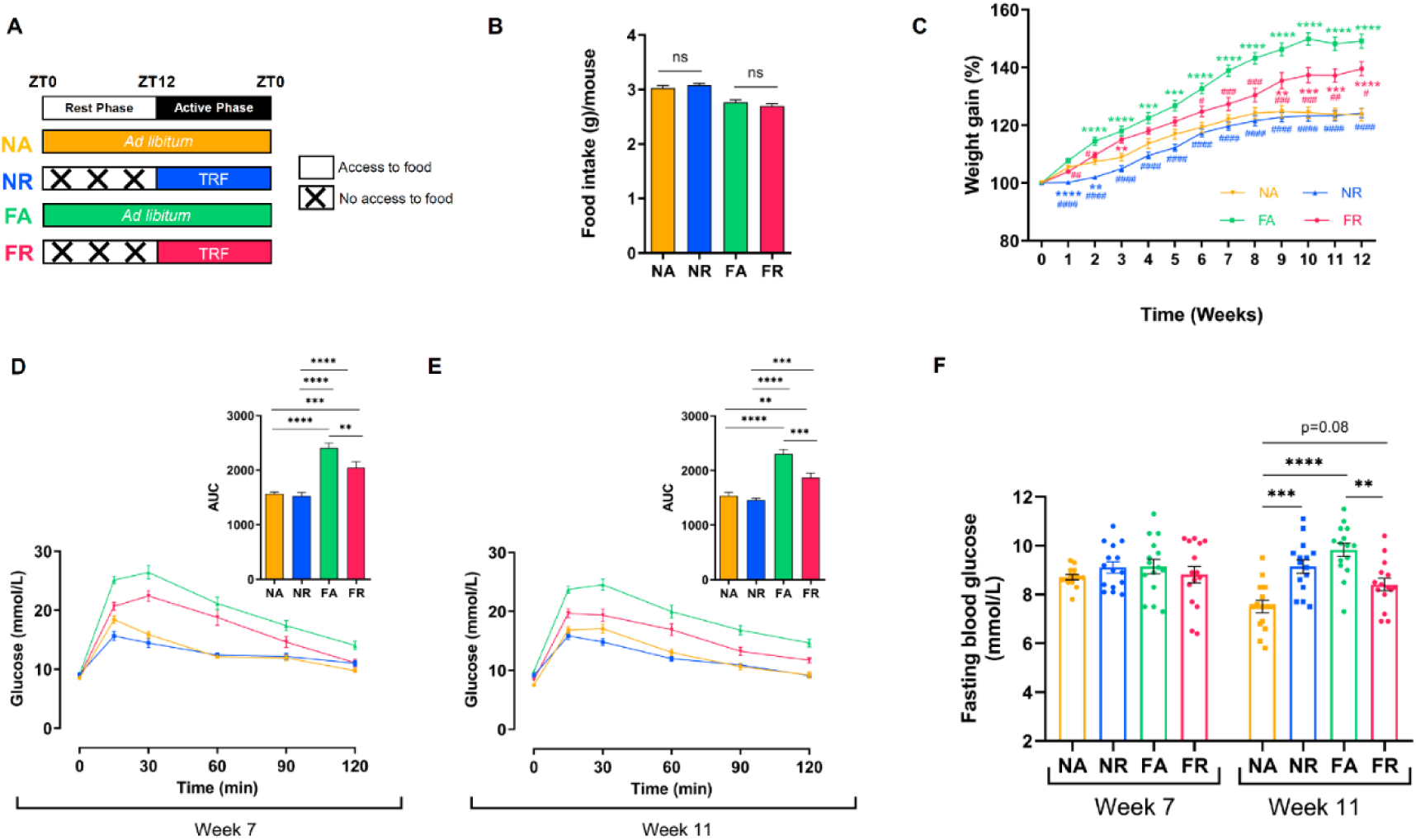
Time-restricted feeding (TRF) mitigated the HFD-induced adverse metabolic effects in mice. (A) Outline of study design indicating the different diet and feeding regimes. NA and FA had *ad libitum* access to food. NR and FR had time-restricted access to food from ZT12 to ZT0. NA and NR were fed a normal-fat diet (NFD), while FA and FR were fed a high-fat diet (HFD). The colored bar indicates access to food. (B) Average daily food intake during the study period. Data expressed as mean ± SEM and analyzed by unpaired *t*-test for each diet. (C) Percent body weight gain over time for each group. Data expressed as mean ± SEM and analyzed by one-way ANOVA followed by Tukey’s test at each time point *p<0.05, **p<0.01, ***p<0.001, ****p<0.0001 (compared to NA); ^#^p<0.05, ^##^p<0.01, ^###^p<0.001, ^####^p<0.0001 (compared to FA). n =15/group except NR, n=14 for week-12. The color of the symbols indicates the group that is significantly different at the specific time point. (D) Oral glucose tolerance test (OGTT) after 7 weeks (n =15/group) and (E) 11 weeks of study duration with the calculated area under the curve (AUC) of blood glucose (n =15/group, except NR, n=14). (F) Fasting blood glucose. (D-F). Data expressed as mean ± SEM and analyzed by one-way ANOVA followed by Tukey’s test at each time point; *p< 0.05, **p < 0.01, ***p < 0.001, ****p < 0.0001.

In general, the NA and NR groups experienced similar weight gain, while TRF reduced weight gain in HFD-fed mice (Fig. 1C). Specifically, the NR group initially showed a significant decrease in body weight gain compared to NA, but the difference between the groups diminished over time, and from week-3 onwards, no significant difference was observed between the two NFD-fed groups. Interestingly, the FR mice gained less body weight than the FA mice, despite having similar food intake. The FR group exhibited significantly lower body weight gain than the FA group during the initial 2 weeks, and then again from week-6 onwards, the difference was statistically significant and remained so until the end of the study. Moreover, the weight gain in the FR group was not significantly different from the NA mice for the first 8 weeks of the study, except for week-3. However, the FA mice gained significantly more weight than the NA mice throughout the entire study.

To determine TRF’s impact on glucose tolerance, an oral glucose tolerance test (OGTT) was performed after 7 and 11 weeks of intervention (Fig.1D-F). The FR mice demonstrated a significant improvement in glucose tolerance, with a faster glucose clearance rate compared to the FA mice during both OGTTs. However, TRF did not affect glucose tolerance in NFD-fed mice, and both groups exhibited similar glucose clearance during the OGTTs. Additionally, fasting blood glucose measured during week-7 showed no significant difference between the groups. Furthermore, fasting blood glucose in the FR mice was significantly lower than in the FA group during the OGTT conducted after 11 weeks of intervention. Interestingly, fasting blood glucose levels measured after 11 weeks of intervention were comparable between the FR and NA mice. In contrast, NA mice had significantly lower fasting blood glucose than NR mice after 11 weeks, but no significant difference in glucose tolerance was observed.

Overall, TRF improved the metabolic outcomes in mice fed with HFD without reducing caloric intake.

### TRF changed the overall gut microbiota structure only in mice fed with HFD

To investigate how the gut microbiota responded to TRF, we sequenced the V3-V4 region of the 16S rRNA gene from the DNA extracted from fecal samples collected from the mice one day before starting the treatment (baseline) and then after 12 weeks of dietary intervention. After sequencing, we obtained 4,864,921 high-quality reads (average 41,228 ± 5,562 reads per sample), denoised into 1,140 amplicon sequence variants (ASVs).

The overall gut microbiota structure was not different between the groups before the treatment began (Fig. 2A and 2B). After 12 weeks of intervention, FR mice exhibited increased gut microbiota diversity (Shannon index) and richness (number of observed ASVs) compared to FA mice (Fig. 2C). However, alpha diversity measures were not significantly different between the NA and NR groups. Principal-coordinate analysis and permutational multivariate analysis of variance (PERMANOVA) based on the Bray-Curtis dissimilarity metric showed that the overall gut microbiota structure in the FA group was significantly different from that in the FR group, but no significant difference was observed between the NA and NR groups (Fig. 2D). This indicates that TRF altered the overall structure of the gut microbiota in HFD-fed mice, consistent with the improvements observed in metabolic outcomes.

**FIG 2.**
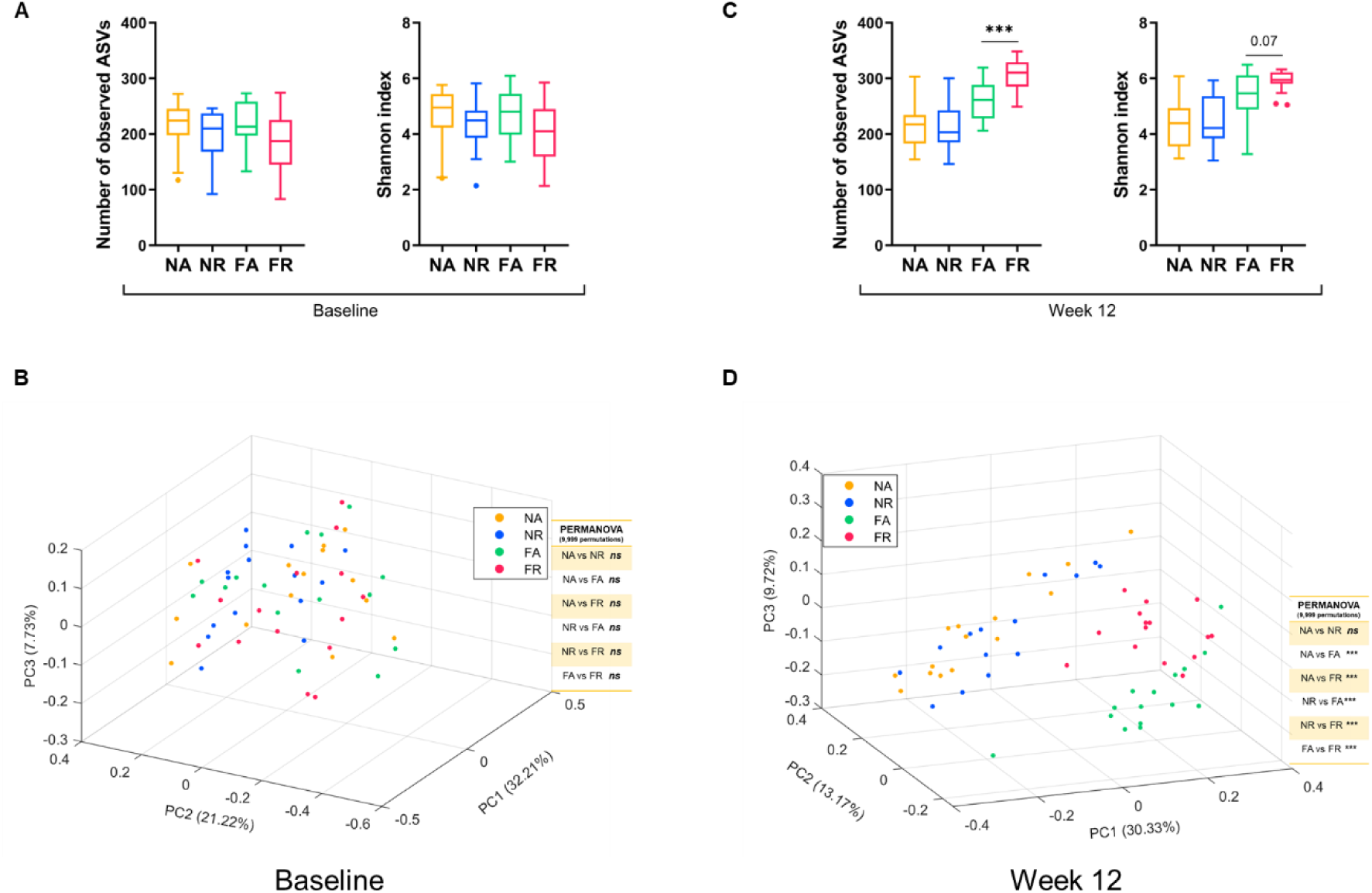
TRF altered the overall gut microbiota structure in HFD-fed mice at the ASV level. Alpha diversity was measured by the number of observed amplicon sequence variants (ASVs) and Shannon index at (A) baseline and (C) after 12 weeks of TRF. Data expressed as mean ± SEM and Mann-Whitney test was used to compare the effect of a difference in feeding regime in each diet. Principal-coordinate analysis (PCoA) plot of the gut microbiota structure based on the Bray-Curtis dissimilarity metric with corresponding PERMANOVA comparisons (9,999 permutations) and p values adjusted with Benjamini-Hochberg correction for multiple group comparison at (B) baseline and (D) after 12 weeks of TRF.

Bacteria in the gut form a network of complex ecological interactions and respond to disturbances as functional groups called guilds. Members within the same guild exhibit co-abundance behavior. We constructed a co-abundance network using 297 ASVs found in over 25% of the samples, accounting for approximately 96% of the total reads, to identify potential guilds that responded to TRF (Fig. 3A and Table S2). These 297 ASVs were clustered into 35 co-abundance groups (CAGs). PCoA analysis performed on the relative abundance profiles for the 35 CAGs using the Bray-Curtis dissimilarity metric revealed that at baseline, the groups had similar gut microbiota structures at the CAG level (Fig. 3B). However, after 12 weeks of dietary intervention, FA and FR groups formed separate clusters in the PCoA plot, while mice from NA and NR groups were indistinguishable (Fig. 3C). Additionally, PERMANOVA analysis demonstrated that FA and FR groups differed significantly, while NA and NR groups did not differ significantly after 12 weeks of intervention (Fig. 3C). Procrustes analysis between the PCoA based on the Bray-Curtis dissimilarity metric for the rarefied ASV dataset and the PCoA on the relative abundance profiles for the 35 CAGs based on the Bray-Curtis dissimilarity metric before and after treatment showed a significant correlation (PROTEST analysis) between the two datasets (Fig. 4). This indicates that the CAG level data provided a good representation of the overall gut microbiota structure that agreed with the ASV level data.

**FIG 3.**
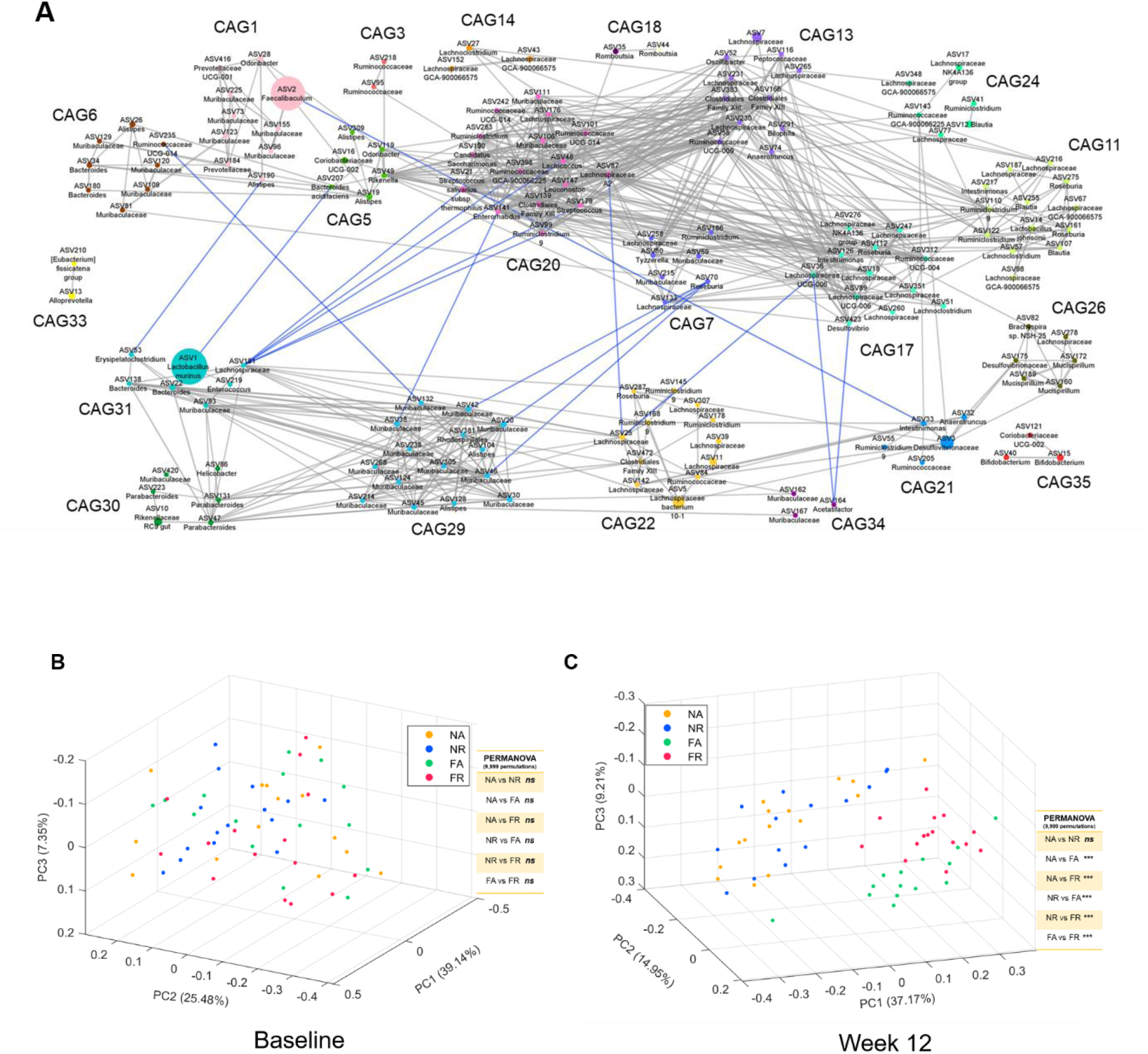
TRF altered the overall gut microbiota structure at the co-abundance group (CAG) level in HFD-fed mice. (A) Microbial interaction network formed with prevalent ASVs from the four groups using baseline and week 12 abundance data. The network plot displayed the interactions between the different CAGs. Node size represents the mean abundance of ASV. The edges between the nodes indicate correlation (gray = positive, blue = negative), with the width of the edge corresponding to the magnitude of the correlation. Correlations with absolute values greater than 0.6 and CAGs with mean abundance greater than 1% are shown here for improved visualization. Principal-coordinate analysis (PCoA) analysis was performed using the relative abundance profiles for the 35 co-abundance groups (CAGs) based on the Bray-Curtis dissimilarity metric at (A) baseline and (B) after 12 weeks of the TRF regime.

**FIG 4.**
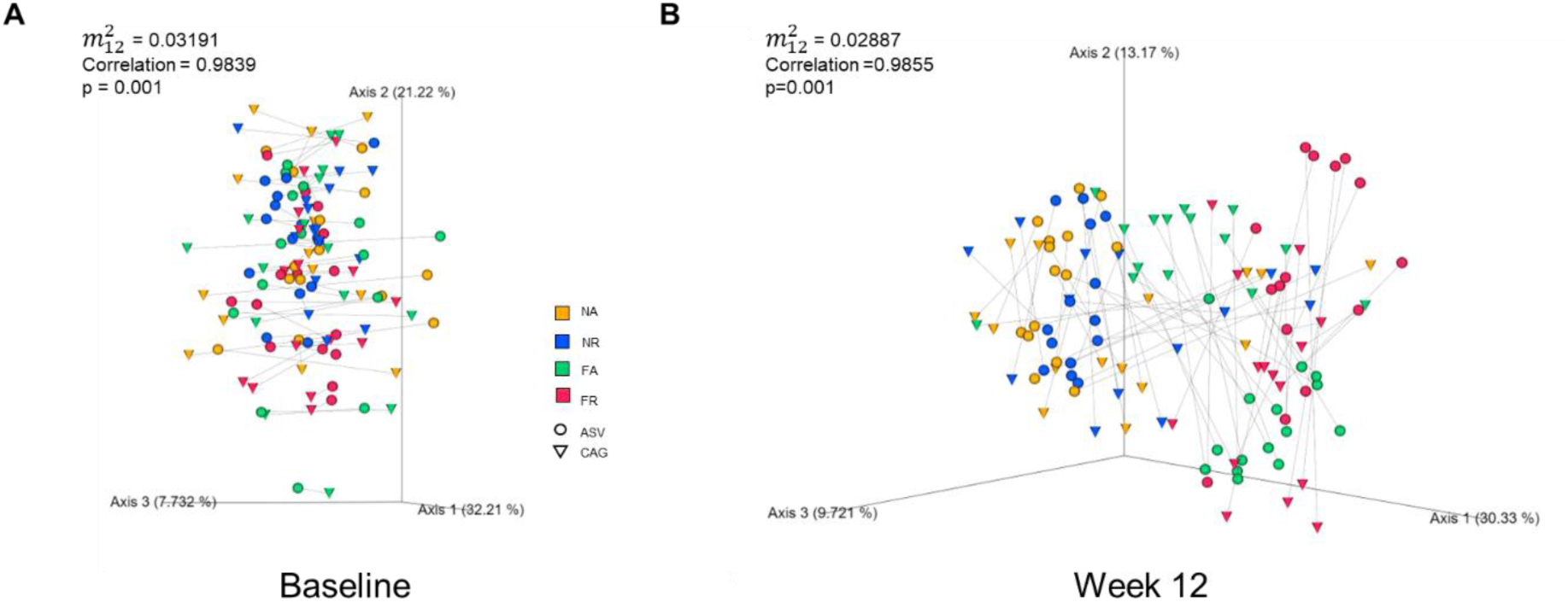
Significant concordance between the gut microbiota structure at ASV and CAG levels. Procrustes analysis was performed on the PCoA based on the Bray-Curtis dissimilarity metric for the rarefied ASV dataset and the PCoA on the relative abundance profiles for the 35 CAGs based on the Bray-Curtis dissimilarity metric at (A) baseline and (B) after implementing 12 weeks of TRF regime. PROTEST analysis was used to test for statistical significance between the two ordinations with 999 permutations.

Next, we conducted a taxon-based analysis at the genus level to compare it with the CAG-level analysis. Out of 1,131 ASVs, only 675 ASVs were annotated at the genus level. These 675 ASVs were classified into 130 genera that accounted for approximately 78% of the total reads. The remaining 456 ASVs, accounting for about 22% of the reads, were excluded from subsequent analysis, as these ASVs unclassifiable at the genus level. PCoA analysis with the Bray-Curtis dissimilarity metric and PERMANOVA test on the genus level relative abundance showed that at baseline, the four groups had similar gut microbiota structures (Fig. S1). However, after 12 weeks of dietary intervention, the FA and FR groups formed two distinct clusters and were significantly different, whereas the NA and NR groups clustered together and did not differ significantly. Procrustes analysis compares two datasets by fitting them together through superimposition and minimizing the sum of squared distances (Procrustes sum of squares, 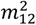) between the positions of the same sample in two different ordination plots. 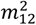 provides a measure of the degree of concordance between the two datasets. Procrustes analysis between the Bray-Curtis dissimilarity metric PCoA plots for the rarefied ASV dataset and the relative abundance profile for the 130 genera before and after treatment showed a significant correlation between the two datasets (Fig. S2). However, 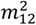 for the genus-level analysis was higher than that for the CAG-level analysis, indicating that CAG-level analysis demonstrates a higher degree of concordance to ASV-level data than genus-level analysis. We calculated an information loss score based on 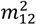, i.e., the ratio between the 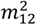 for genus and CAG-level analysis. At baseline, the information loss score was calculated as 5.15, and after 12 weeks of intervention, it was 1.50. Moreover, the information loss scores for the 297 ASVs (grouped into 35 CAGs) vs. 1,131 ASVs and 675 ASVs (classified into 130 genera) vs. 1,131 ASVs were 5,094.5 and 11.13 for baseline and after 12 weeks of intervention, respectively. This suggests that CAG-level analysis preserves the overall gut microbiota structure and prevents information loss better than genus-level analysis.

In summary, TRF altered the overall gut microbiota structure in HFD-fed mice, which was consistent with the improved metabolic phenotype observed in these mice. Furthermore, the CAG level data (35 CAGs) preserved the overall gut microbiota structure observed at the ASV level (1,131 ASVs) better than the taxon-based analysis. This suggests that CAG analysis can be used to reduce the high dimensionality and sparsity associated with ASV-level data.

### TRF-induced alterations in gut microbiota were associated with improved metabolic outcomes in HFD-fed mice

TRF differentially altered the abundance of CAGs in the FA and FR groups. We used Boruta, a Random Forest-based feature selection algorithm, to identify the CAGs that differentiated between the FA and FR groups. We found a set of 11 CAGs that could distinguish between FA and FR mice after 12 weeks of intervention (Fig. 5A). The area under the ROC curve (AUC-ROC=0.95) (Fig. 5B) was used to assess the performance of the classification model. Six out of the 11 CAGs identified were significantly correlated with metabolic outcomes. Among these, five CAGs—CAG3, CAG11, CAG12, CAG31, and CAG33—were significantly enriched in FR mice (Fig. 5D and E). Moreover, these five CAGs were significantly associated with improvements in metabolic outcomes (Fig. 5C). CAG3, CAG12, and CAG33 negatively correlated with glucose intolerance, while CAG11, CAG12, and CAG31 negatively correlated with fasting blood glucose. CAG31, with the dominant ASV from Lactobacillus murinus, was significantly increased in FR mice compared to the FA group. Also, CAG11, with ASVs from *Lactobacillus johnsoni*, *Lachnoclostridium*, Lachnospiraceae, *Blautia*, *Roseburia*, and *Ruminiclostridium*, was significantly promoted in FR mice. CAG3, increased in FR mice, primarily comprised ASVs from Lachnospiraceae and Ruminococcaceae families. Again, CAG12 showed increased abundance in FR mice with ASVs from Lachnoclostridium, Roseburia, Lachnospiraceae, and Ruminococcaceae families. Additionally, CAG33 with predominant ASVs from the *Alloprevotella* genus was increased in FR mice. Furthermore, CAG1 was positively associated with the deterioration of glucose tolerance and fasting blood glucose (Fig. 5C). The relative abundance of CAG1 was significantly increased in FA mice. CAG1 mainly consisted of ASVs from *Fecalibaculum*, *Odoribacter*, and Muribaculaceae.

**FIG 5.**
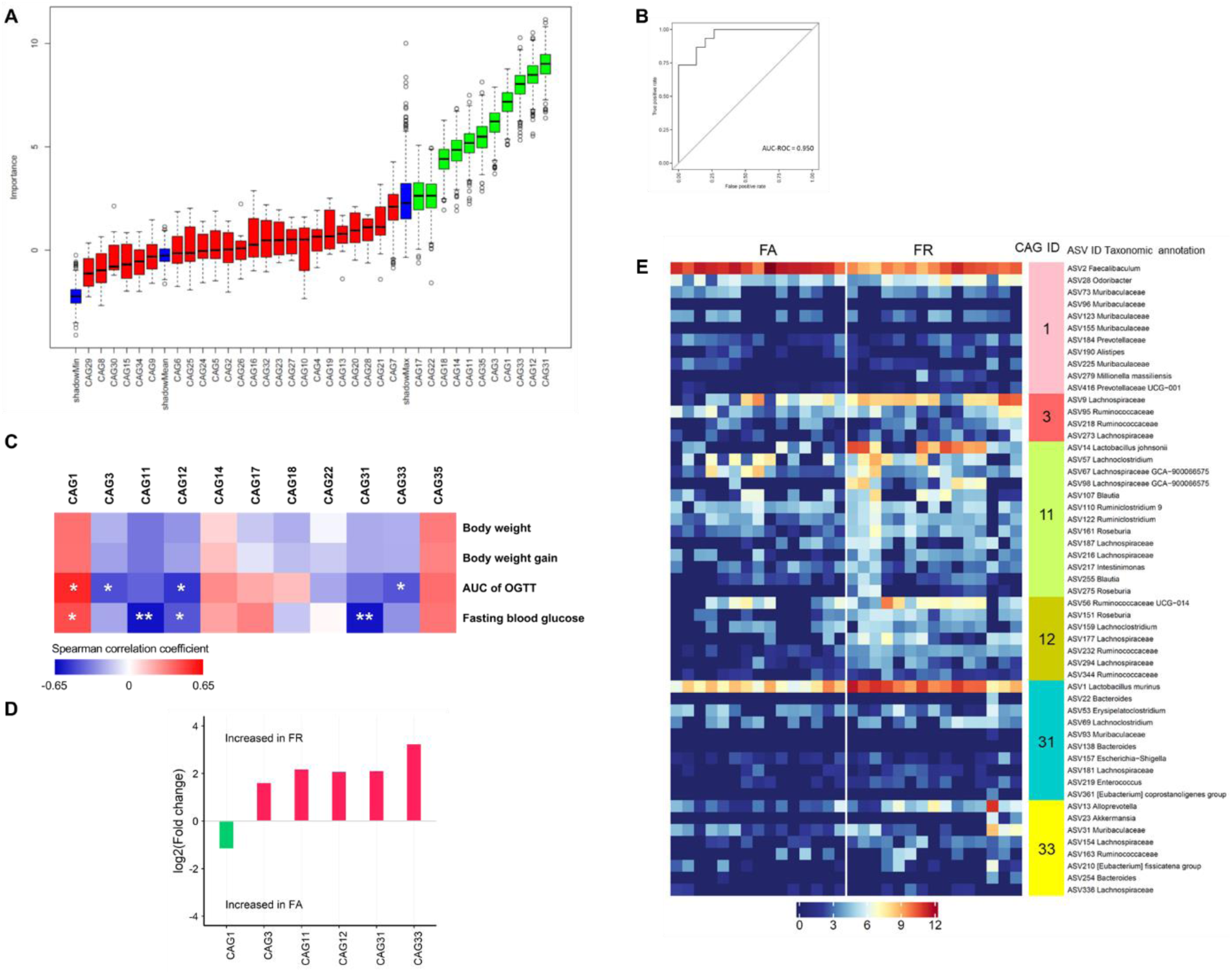
TRF-induced CAGs were associated with improved metabolic outcomes in HFD-fed mice. TRF-induced CAGs were associated with metabolic outcomes in HFD-fed mice. (A) Boxplot of the importance score (Z score) of the CAGs identified by Boruta for differentiating between FA and FR after 12 weeks of intervention. The boxplots in “green” were identified as key variables capable of discriminating between FA and FR, whereas the boxplots in “red” were found as nondiscriminatory. The boxplots in “blue” refer to the minimum, mean, and maximum Z scores of a shadow variable. (B) The area under the ROC curve for the CAG-based classification of the FA and FR group (AUC=0.950; 95% CI: 0.87,1.03); ROC: Receiver operating characteristic. (C) Heatmap of Spearman’s correlation (with FDR correction) between the relative abundance of the discriminating CAGs identified by Boruta, and the metabolic parameters related to glucose and lipid metabolism. *p< 0.05, **p < 0.01, ***p < 0.001. (D) Barplot of the Log2 fold change in the relative abundance of the key CAGs and (E) heatmap of relative abundance of the ASVs (log2-transformed) which form the key CAGs that showed significant correlation with the host metabolic parameters.

In summary, TRF altered the gut microbiota composition in HFD-fed mice. TRF-induced specific alterations in the gut microbiota composition were associated with improvements in metabolic outcomes in HFD-fed mice.

### TRF restored diurnal rhythmicity in the overall structure of the gut microbiota

To determine the effect of TRF on the cyclical dynamics of the gut microbiota in HFD and NFD-fed mice, we analyzed the gut microbiota structure after 11 weeks of the scheduled feeding regime by collecting fecal samples every 6 hours over two consecutive light-dark cycles. The DNA was extracted from the fecal samples, and the V3-V4 region of the 16S rRNA gene was sequenced. A total of 3,706,340 high-quality reads (average 30,886 ± 5365.73) were obtained and further denoised into 917 ASVs. The ASV level data were reassembled into the 35 CAGs that were constructed earlier. We identified 295 out of the 297 prevalent ASVs that were previously used to construct the CAGs in the week 11 microbiota data. These 295 ASVs accounted for ~93% of the total number of reads. We used the empirical JTK_CYCLE algorithm to detect rhythmicity in the microbial sequencing data at the CAG level. The PCoA analysis revealed a robust diurnal pattern in the overall microbial community structure in the FR and NR mice compared to their counterparts (Fig. 6A and B, S3A and B). Consistently, the respiratory exchange ratio (RER) and energy expenditure (EE) from metabolic cage recordings also showed improved rhythmic patterns in the TRF groups (Fig. 7).

**FIG 6.**
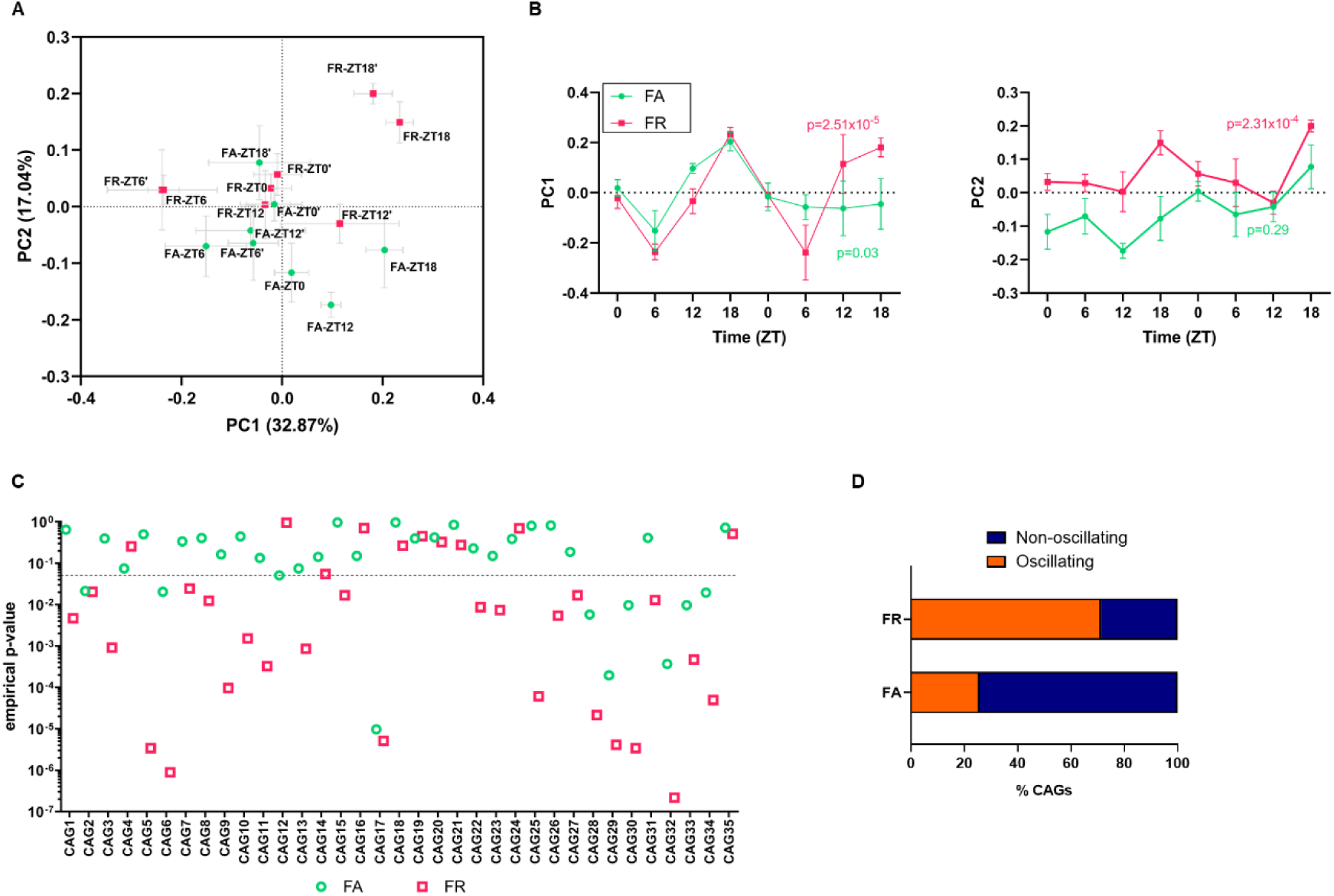
Time-restricted feeding (TRF) displayed robust diurnal rhythmicity in gut microbiota’s overall structure and composition at the CAG level compared to mice with *ad libitum access to* HFD. (A) PCoA analysis was performed at the CAG level using the abundance profile of the 35 CAGs based on the Bray-Curtis dissimilarity of all the samples collected throughout two light-dark cycles during week-11 of treatment in HFD-fed mice. (B) Alteration of the gut microbiota structure along the first and second principal coordinate (PC1 and PC2) of the PCoA based on Bray-Curtis dissimilarity in HFD-fed mice. Data were plotted as mean ± SEM.; n=3-5 fecal samples per time point. Rhythmicity was analyzed using the non-parametric empirical JTK_CYCLE algorithm and p-values refer to empirical-p values. (C) CAGs display diurnal oscillation in their relative abundance under TRF and *ad libitum* feeding regimes in HFD-fed mice. Rhythmicity was examined using the non-parametric empirical JTK_CYCLE algorithm and p-values refer to empirical-p values. The dashed line indicates p<0.05. (D) Percentage of oscillating and non-oscillating CAGs.

**FIG 7.**
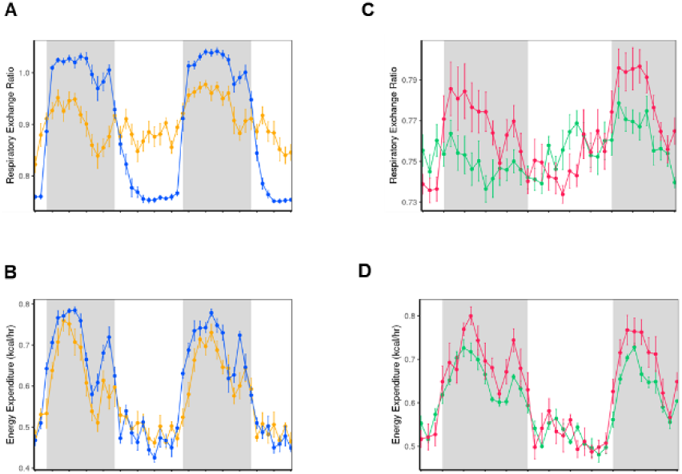
Respiratory exchange ratio (RER) and energy expenditure (EE) from metabolic cages recordings in (A, B) NFD-fed mice and (C, D) HFD-fed mice after 12 weeks of dietary intervention (n=5/group). The gray color indicates the active phase, and the white color indicates the rest phase.

Next, we determined whether the proportion of CAGs undergoing cyclical oscillation in their relative abundance varied between the feeding regimes. As the NA group did not display diurnal rhythmicity in either PC1 or PC2 it was not included in further comparisons. FR mice displayed a higher percentage of cyclical CAGs than the FA group (Fig. 6C and D, Table S4). NR group also displayed oscillation in 3 CAGs but much lower than the FR group (Fig. S3C, Table S4). TRF induced a substantial increase in the proportion of CAGs undergoing diurnal oscillation in the HFD-fed mice, from 9 CAGs in FA to 25 in FR (Fig. S4). Thus, TRF introduced a significant change in the overall cyclical dynamics of the gut microbiota in the HFD-fed mice compared to the NFD-fed mice. To investigate whether the CAGs associated with improved host metabolic outcomes exhibited diurnal variation in their relative abundance under the TRF regime in HFD-fed mice, we applied the empirical JTK_CYCLE algorithm to the CAG relative abundance data. Four out of the five CAGs (CAG3, CAG11, CAG31, and CAG33) that increased in the FR group displayed significant diurnal oscillation in their relative abundance compared to the FA group (Fig. 8). CAG12 did not oscillate in its relative abundance; instead, it remained stable throughout the 48 hours. Moreover, the relative abundance of CAG12 was higher in the FR group than in the FA group over the 48-hour duration. Additionally, CAG1, which was associated with impaired glucose metabolism in mice, displayed significant diurnal oscillation in the FR group. However, CAG1 remained relatively constant with increased relative abundance in the FA group compared to the FR group throughout the 48 hours. Thus, TRF restored diurnal rhythmicity in most of the key CAGs associated with glucose metabolism in HFD-fed mice.

**FIG 8.**
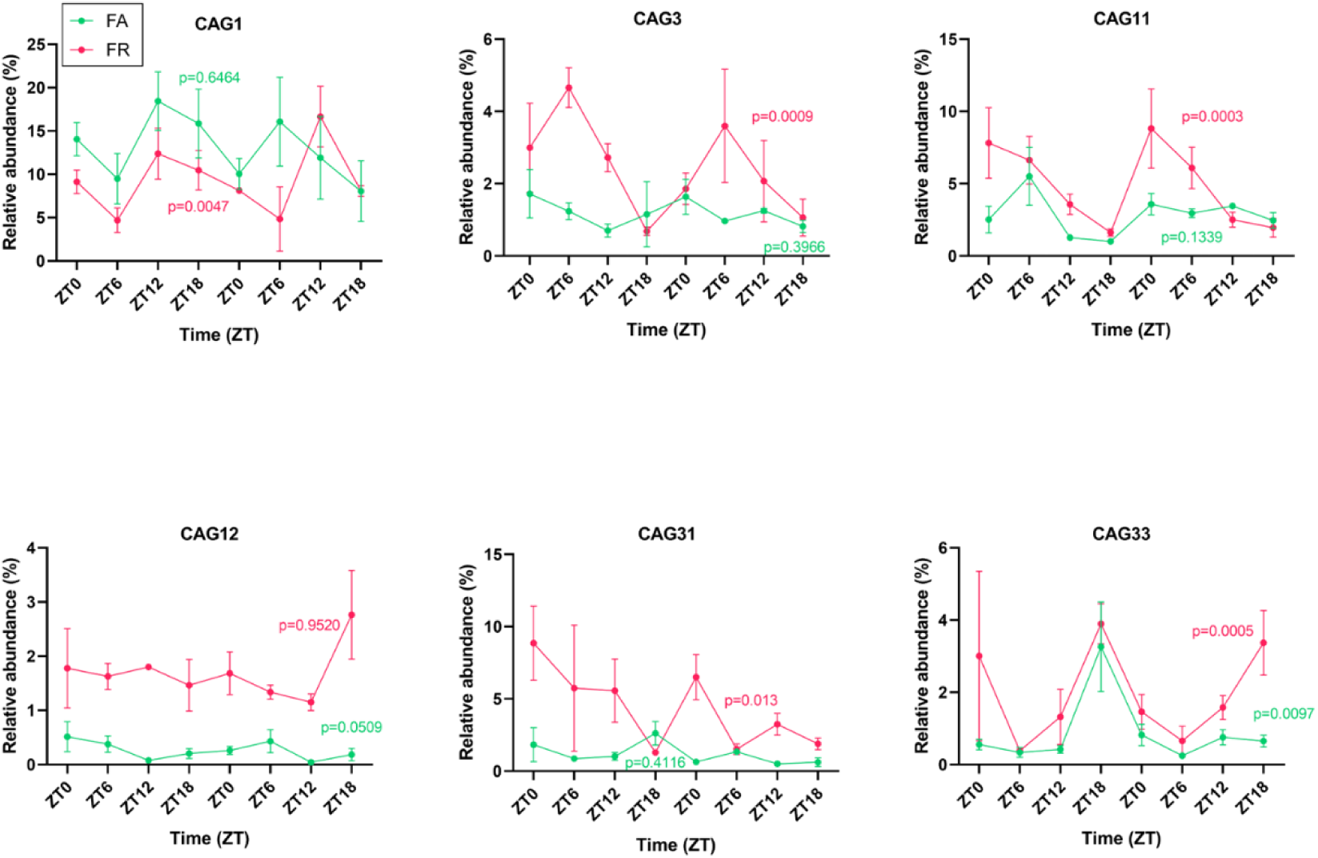
Key CAGs associated with host metabolic parameters in HFD-fed mice displayed differences in their diurnal pattern. Data were plotted as mean ± SEM; n=3-5 fecal samples per time point. Rhythmicity was determined using the non-parametric empirical JTK_CYCLE algorithm and p-values refer to empirical-p values.

In summary, TRF promoted diurnal oscillation in the overall structure of the gut microbiota in HFD-fed mice. Furthermore, most of the key CAGs associated with improved metabolic outcomes in HFD-fed mice regained diurnal rhythmicity under the TRF regime. This suggests that the imposed diurnal rhythm in the gut microbiota may contribute to the improved metabolic phenotype observed in HFD-fed mice.

## Discussion

In this study, we discovered that time-restricted feeding (TRF) significantly enhanced body weight gain and glucose intolerance in high-fat diet (HFD)-fed mice without lowering caloric intake. These metabolic health benefits were linked to selective changes in the gut microbiota at the guild level. This finding is especially noteworthy because it shows that consuming the same amount and type of food, but only during the active phase, can substantially modify the gut microbiota in HFD-fed mice, leading to improved body weight gain and glucose tolerance. This suggests that the mitigation of HFD-induced metabolic dysfunction might be mediated by key guilds that exhibit different responses to active-phase feeding and associations with glucose homeostasis. Additionally, TRF encouraged diurnal oscillations in most of the essential co-abundance groups (CAGs) that responded to the TRF regimen and were linked with glucose metabolism in the HFD-fed mice. The restoration of diurnal oscillations in most key guilds in response to the feeding-fasting cycle in time-restricted feeding may play a critical role in alleviating metabolic deterioration in ad libitum HFD feeding.

We demonstrated that the guild-based analysis preserved the information provided by the amplicon sequence variant (ASV) level data more effectively than genus-level taxon-based analysis. Potential guilds (CAGs) were assembled by clustering individual members based on the co-abundance patterns of prevalent ASVs. Furthermore, a significantly higher percentage of sequencing reads were retained in the guild-based analysis compared to the genus-level analysis. Since many gut ecosystem members are novel, excluding members that cannot be annotated to a specific taxonomic level for analysis may result in misleading interpretations. For instance, ASV9 could only be assigned to the family level (Lachnospiraceae), and this ASV was removed from the taxon-based analysis because a genus-level annotation was unavailable (Table S2). However, in guild-based analysis, ASV9 was part of CAG3, which was negatively correlated with phenotypic outcomes. Unlike taxon-level analysis, where different members of a taxon function in unrelated ways, the guild-based approach offers an ecologically relevant tool for identifying key gut microbial community members associated with a specific host phenotype. This method thus presents an unsupervised, reference database-independent technique based on ASV co-abundance (in conjunction with a longitudinal study design) for identifying potential guilds while preserving the information conveyed by ASV-level data.

In the current study, TRF-induced changes in six CAGs were associated with metabolic outcomes in HFD-fed mice. CAG3, CAG11, CAG12, CAG31, and CAG33 were linked to improved glucose metabolism and selectively promoted in the FR group compared to the FA mice. CAG31 contained the dominant ASV1 from Lactobacillus murinus, and the increased abundance of L. murinus has previously been shown to correlate with improved metabolic phenotype in mice^22^. ASV14 in CAG11 was identified as a member of L. johnsonii. Several strains of L. johnsonii have demonstrated the ability to inhibit pathogenic bacteria in the gut, exhibit acid resistance properties, and show anti-inflammatory effects in mice^23,24^. It is suggested that the ferulic acid esterase activity displayed by some L. johnsonii strains can plausibly improve diabetes, as ferulic acid has been shown to stimulate insulin secretion in rodents^25^. Some members of the remaining CAGs also showed associations with improved glucose metabolism, such as Lachnoclostridium, Blautia, Roseburia, and other Lachnospiraceae family members. Members of the Lachnospiraceae family have been reported to produce short-chain fatty acids (SCFAs) via fermentation of various complex carbohydrates, which in turn contribute to the beneficial effects observed in metabolic diseases or produce bacteriocins to potentially inhibit the growth of other bacteria^26,27^. Although bacteria from the Lachnospiraceae family have also been implicated in metabolic dysfunction^26^, ASVs in this family were in the CAGs associated with improved metabolic health, suggesting that these Lachnospiraceae family members were part of a beneficial guild. Additionally, ASV13 in CAG33 was identified as a member of the Alloprevotella genus, which can produce acetic and succinic acid^28^, reported to improve obesity-induced metabolic impairments^29,30^. On the other hand, CAG1, consisting mostly of ASVs from Faecalibaculum and Muribaculaceae, was associated with higher fasting glycemic levels and impaired glucose tolerance, and CAG1 was increased in the FA group compared to the FR group. This indicates that CAG1 may be a guild of pathobionts. Previously, the *Faecalibaculum* genus was associated with increased body weight gain, glucose intolerance, and serum lipopolysaccharide-binding protein^31^. Members of the Muribaculaceae family (previously classified as Bacteroidales S24-7 family) have not been well studied; it has been reported that some of the family members can degrade mucin and were promoted by low-level inflammation in mice^32,33^. ASVs assigned to Muribaculaceae in CAG1 were potentially pathogenic. The guild-based analysis allowed us to cluster known or unknown bacteria into the same guild and helped identify novel beneficial or pathogenic bacteria based on prior knowledge about the members in the same guild.

Previous studies have shown that the gut microbiota composition and function undergo diurnal fluctuations, and these daily rhythms are strongly influenced by feeding time and the macronutrient composition of the diet^7–9^. Genetic disruption of the circadian clock disturbs the cyclical oscillation in gut microbiota composition, which can be partly restored by timed feeding^7,34^. In this study, we demonstrated that the TRF regime promotes daily oscillations in the overall structure and composition of the gut microbiota at the guild level. Furthermore, we found that most of the key CAGs associated with improved glycemic control in HFD-fed mice showed enhanced diurnal rhythm in the TRF group. This suggests that not only the differential abundance of the key guilds but also the diurnal oscillatory behavior of these guilds may be an important factor in mediating the beneficial effects of TRF in HFD-fed mice.

Interestingly, the CAGs containing bacteria from the *Lactobacillus* genus (CAG31 and CAG11) showed cyclical oscillation in the FR group, as observed previously^20^. TRF-induced increased cyclical oscillation in gut microbiota composition was consistent with the strength of improvement in metabolic outcomes. Moreover, a recent study performed in a human cohort showed that type 2 diabetes was associated with disrupted rhythmicity in gut microbiota composition and identified a set of arrhythmic bacterial taxa that could predict diabetes^35^. However, the mechanisms linking these diurnal microbial oscillations and metabolic health need further study. Niche modification, in which the activity of the initial set of gut bacteria alters their local environment, creating new niches to support the growth of other bacteria, may drive the diurnal dynamics of the gut microbiota^36^. For example, in this study, CAG 31 showed significant oscillation in the FR group but not in the FA group. Non-oscillating microbes, on the other hand, do not respond to the changing gut environment and are relatively stable throughout the day. For example, CAG12 did not show an oscillatory pattern but its relative abundance was significantly higher in FR than in FA. Furthermore, TRF-induced cyclical oscillations in the gut microbiota might help restore the cyclical oscillation in microbially-derived metabolites, such as SCFAs, which may affect host metabolism. It has been reported that butyrate displays a robust diurnal rhythm in cecal and fecal samples collected from mice fed a regular chow diet ad libitum, while the rhythmicity in butyrate was lost when mice were fed an HFD^9^. SCFAs can also modulate host peripheral clock gene expression in both in vitro and in vivo systems^9,37^. Thus, SCFA-producing guilds such as CAG11 and CAG31 may modulate host metabolic rhythmicity with a diurnal rhythm of their abundance in the gut.

TRF restored diurnal rhythmicity of the overall structure of the gut microbiota at the CAG-level in both HFD and NFD-fed mice. But a greater number of CAGs regained diurnal oscillation in the FR group compared to the NR group. For the ad libitum-fed groups, NA did not show diurnal behavior in the structure of gut microbiota (CAG-level), and FA displayed weak diurnal oscillatory behavior (CAG-level), which is in contrast with previous studies^8,9^. Such differences may arise due to the composition of the control diets used in this study. In prior studies^8,9^, the HFD used was made with purified ingredients, whereas the control diet was a grain-based diet with high amounts of dietary fiber from varied sources^38–40^. As dietary fiber is a critical ingredient that shapes the gut microbiota^38,39,41^, it may impact the diurnal rhythm observed in the gut microbiota. Additionally, dietary fiber content can influence gut transit time^42,43^ and SCFAs production^39^, which can potentially affect the oscillatory behavior of the gut microbiota. As previously reported, the composition of the diet can alter gut motility; for example, HFD can increase gut transit compared to a control diet^42,43^. Thus, in the current study, HFD and NFD were prepared similarly with purified ingredients and a similar amount of dietary fiber to prevent confounding effects. However, further studies are needed to understand the impact of dietary fiber and fat on the diurnal dynamics of the gut microbiota.

In summary, our study demonstrated that TRF can improve HFD-induced body weight gain and glucose intolerance in association with specific alterations in the structure of the gut microbiota at the guild level, even without reductions in calorie intake. Moreover, most of the responding bacterial guilds displayed diurnal oscillating patterns under the TRF regime in HFD-fed mice. The alterations in the overall structure and diurnal oscillations of key guilds in the gut microbiota induced by TRF may, in part, contribute to reducing the adverse metabolic effects of HFD. Further research is needed to understand the mechanisms underlying how members of the TRF-enriched guilds of bacteria help mediate beneficial metabolic responses in the host.

## Materials and methods

### Animal experiment and sample collection

All the animal experiments were approved by the Institutional Animal Care and Use Committee (IACUC) of the School of Life Sciences and Biotechnology of Shanghai Jiao Tong University (No. 2018035).

Specific pathogen-free (SPF), 5-week-old male mice (C57BL6/J) were purchased from SLAC Inc. (Shanghai, China) and allowed to acclimatize at the animal center of Shanghai Jiao Tong University for 3 weeks before the start of the experiment. All mice were kept under a strict 12h light: 12h dark cycle, with lights being turned on from 7 a.m. to 7 p.m. (Zeitgeber time: ZT 0 denotes lights on and ZT12 denotes light off) and a constant temperature of 22° C ±3°C. During the acclimation phase, the mice had *ad libitum* access to a normal-fat diet (NFD, Research Diets D12450J; 70% carbohydrate, 20% protein, 10% fat, 3.85 kcal/g; Table 3.1) and autoclaved water. After the acclimation phase the mice were randomly assigned to one of the following four groups (i.e., 15 mice per group and 3 mice per cage) for 12 weeks as described in Fig. 2.1A (i) fed NFD with *ad libitum* access (NA) (ii) fed NFD with access to food restricted to a 12h period between 7 p.m. to 7 a.m. (NR) (iii) fed HFD with *ad libitum* access (FA; HFD, Research Diets D12492; 60% fat, 20% protein, and 20% carbohydrate, 5.21 kcal/g; Table 2.1), and (iv) fed HFD with access to food restricted to a 12h period between 7 p.m. to 7 a.m. During the study period, access to food was regulated by transferring the mice daily between cages that had food and water (feeding-cage) to cages with only water (fasting-cage) for time-restricted feeding groups (NR and FR). To control for the effect of transferring the mice between cages in the time-restricted feeding group, the mice with *ad libitum* access to food were also transferred between two feeding cages at the same time. The body weight of the individual mice and food intake per cage was measured twice weekly at the same time every week. At the end of the study period, mice were sacrificed after 6 hours of fasting (ZT0 to ZT6) followed by a collection of serum and tissue samples, which were stored at −80°C until further analysis. Fresh fecal samples (2-3 fecal pellets/mice) were collected from the mice individually and stored at −80°C until DNA extraction. At the end of 12 weeks of dietary intervention, indirect calorimetry was performed on a subset of mice (n=5/group) using a Comprehensive Lab Animal Monitoring System (CLAMS, Columbus Instruments) at the Shanghai Model Organisms Center, Inc., (Shanghai, China). Mice were placed individually in the metabolic chambers for three consecutive days including 24 hours of acclimatization duration. Light, temperature, and feeding conditions were maintained the same as in the home cages.

### Oral glucose tolerance test

Baseline blood glucose was measured after 6 hours of fasting (ZT0 to ZT6) with the help of a blood glucose meter (ACCUCHEK® Performa, Roche, USA). Glucose (2.0 g/kg body weight) was administered by oral gavage and glucose concentration was determined in the blood collected from the tip of the tail vein at 15, 30, 60, 90, and 120 min.

### Microbial 16S rRNA gene V3-V4 region sequencing and data analysis

Microbial DNA was extracted from the fecal samples collected from the individual mice on the day before the start of the feeding regime for baseline samples and subsequently after 12 weeks of the feeding regime to explore the variation of gut microbiota during the study period, according to a method described previously^44^. To reduce variation from the diurnal oscillating nature of the gut microbiota^20,45^, fecal samples for the baseline and after 12 weeks of dietary intervention were collected at the same time at ZT1 (8 a.m.). During week 11 of the study period, fecal samples from the individual mice were collected every 6 hours over 48 hours (n=3-5/group). The hypervariable V3-V4 region of the 16S rRNA gene was sequenced using the MiSeq sequencing platform (Illumina Inc., USA). A sequencing library for the V3-V4 regions of the 16S rRNA gene was prepared according to a modified version of the instructions provided by the manufacturer (Part # 15044223 Rev. B; Illumina Inc., USA) with some modifications^46^ and sequenced using the Illumina Miseq System (Illumina Inc., United States).

The 16S rRNA gene amplicon sequence data were processed and analyzed on the QIIME2 software (v2019.7)^47^. The raw sequence data were demultiplexed and then denoised with the DADA2 pipeline (q2-dada2 plugin)^48^ to obtain the amplicon sequence variants (ASVs) frequency data table. Alpha diversity metrics (Observed ASVs and Shannon’s index), beta diversity metric (Bray-Curtis dissimilarity), and Principal Coordinate Analysis (PCoA) were performed using the “core-metrics-phylogenetic” plugin after rarefying the samples to 31,500 sequences per sample for baseline and week 12 data. R “ggplot2” package and MATLAB (R2019b) were used to plot the graphs. The alpha diversity indices were compared with Mann-Whitney tests. Statistical significance between the groups was assessed by permutational multivariate analysis of variance (PERMANOVA) with 9,999 permutations and p values obtained were adjusted with the Benjamini-Hochberg correction method^59^. Taxonomic assignment for ASVs was performed via the q2-feature-classifier^49^ using the SILVA 132 16S rRNA gene database^50^. For the week 11 data, the samples were rarefied to 18,000 per sample for subsequent analysis. Beta diversity metric (Bray-Curtis dissimilarity) and PCoA analysis were computed as mentioned earlier in this section. Diurnal oscillatory patterns in principal components/CAGs/ASVs were detected in the fecal samples collected over 48 hours using empirical JTK_CYCLE^51^ with a set period of 24 hours^52^.

### Microbial CAG network analysis

The correlation coefficient between individual ASVs that were present in more than 25% of the samples was calculated using the method described by Bland and Altman for repeated observations^53,54^. The correlation coefficient matrix was converted into a distance matrix (1-correlation coefficient) and subsequently clustered by applying Ward’s hierarchical clustering method to build a tree. Then, PERMANOVA (9,999 permutations) was applied consecutively from the top of the tree to determine distinct clades of the tree as individuals CAGs using a cutoff of p<0.001. The CAG network was visualized in Cytoscape v3.8.2^55^. PCoA analysis based on Bray-Curtis dissimilarity for the formed CAGs and PERMANOVA (9,999 permutations) test to identify statistical significance between the groups was done using QIIME2. Procrustes analysis was then performed between the PCoA of CAGs and the PCoA of ASV data to identify concordance between the two datasets using the “procrustes-analysis” plugin on QIIME2. PROTEST from the R “vegan” package was used to assess statistical significance between the two datasets. Boruta, a Random Forest-based feature selection method, was used to identify the most important CAGs capable of discriminating between the groups^56^. The receiver operating characteristic (ROC) curve was generated using the “evalm” function from the R package “MLeval” for validation of the CAG-based classification.

### Statistical analysis

Statistical analysis was performed using Graphpad Prism 8 software and R (version 3.6.2). The following R packages were used for analysis; “ComplexHeatmap”^57^ and “ggplot2”. One-way ANOVA followed by a Tukey *post hoc* test was used to assess the statistical significance of the physiological and biochemical data. Significance was determined as a p-value < 0.05. The data obtained from the metabolic cage was analyzed with CalR software^58^. Spearman correlation analysis was used to determine the association between CAGs, and metabolic phenotypes followed by FDR correction using Benjamini and Hochberg^59^.

## Data availability

The 16S rRNA gene sequence data generated in this study has been submitted to Sequence Read Archive (SRA) maintained by NCBI under the accession number PRJNA975214.

