## Supplemental Figure 1 for "Guild-level response of the gut microbiota to isocaloric time-restricted feeding in high-fat diet-fed mice"

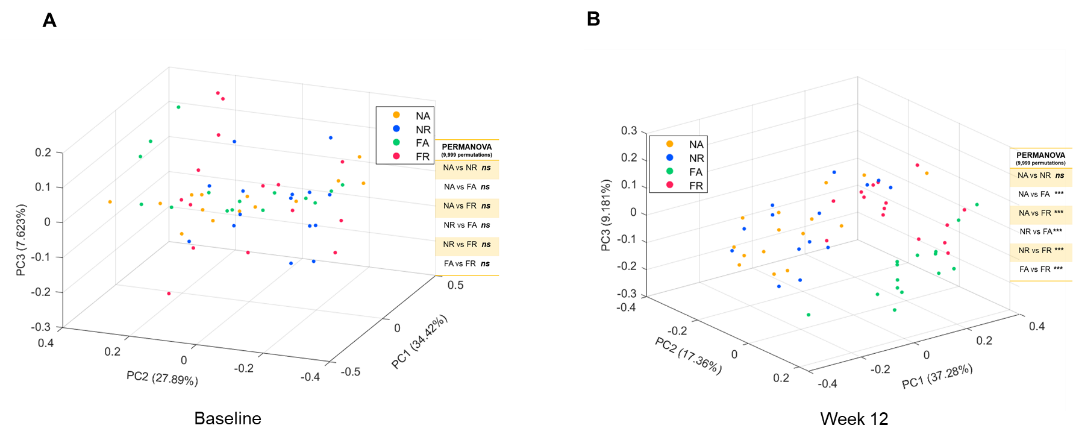

FIG S1	TRF altered the gut microbiota structure at the genus-level in HFD-fed mice. Principal-coordinate analysis (PCoA) was performed using the relative abundance profile at the genus-level with 130 genera based on the Bray-Curtis dissimilarity metric at (A) baseline and (B) after 12 weeks of TRF regime.
