## Supplemental Figure 2 for "Guild-level response of the gut microbiota to isocaloric time-restricted feeding in high-fat diet-fed mice"

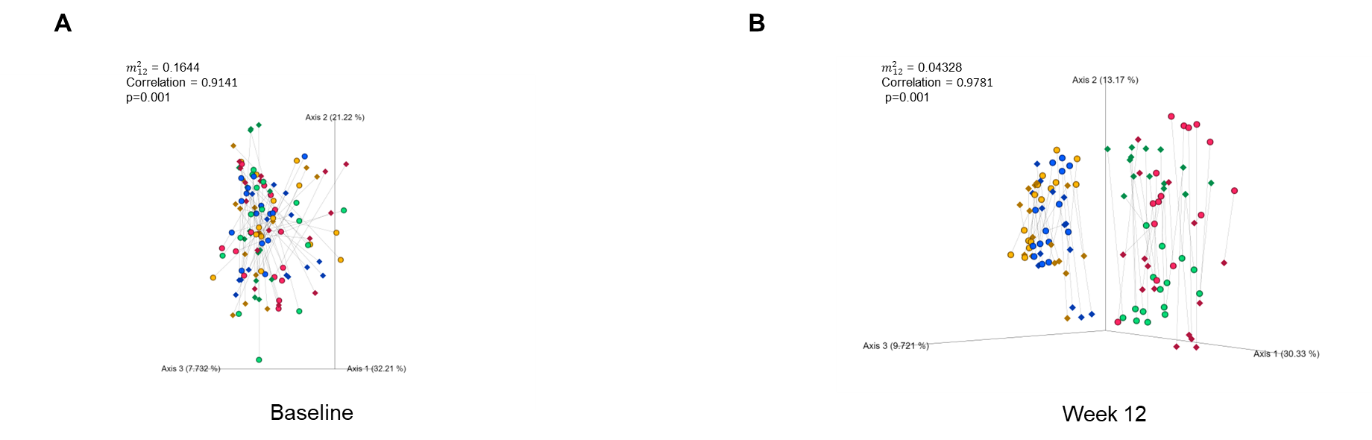

FIG S2	Concordance between the gut microbiota structure ay ASV and genus levels. Procrustes analysis was performed on the Bray-Curtis dissimilarity PCoA plots derived from the rarefied ASV dataset and the relative abundance profiles for the 130 genera at (C) baseline and (D) after implementing 12 weeks of TRF regime. PROTEST analysis was used to test for statistical significance between the two ordinations with 999 permutations.
