## Supplemental Figure 3 for "Guild-level response of the gut microbiota to isocaloric time-restricted feeding in high-fat diet-fed mice"

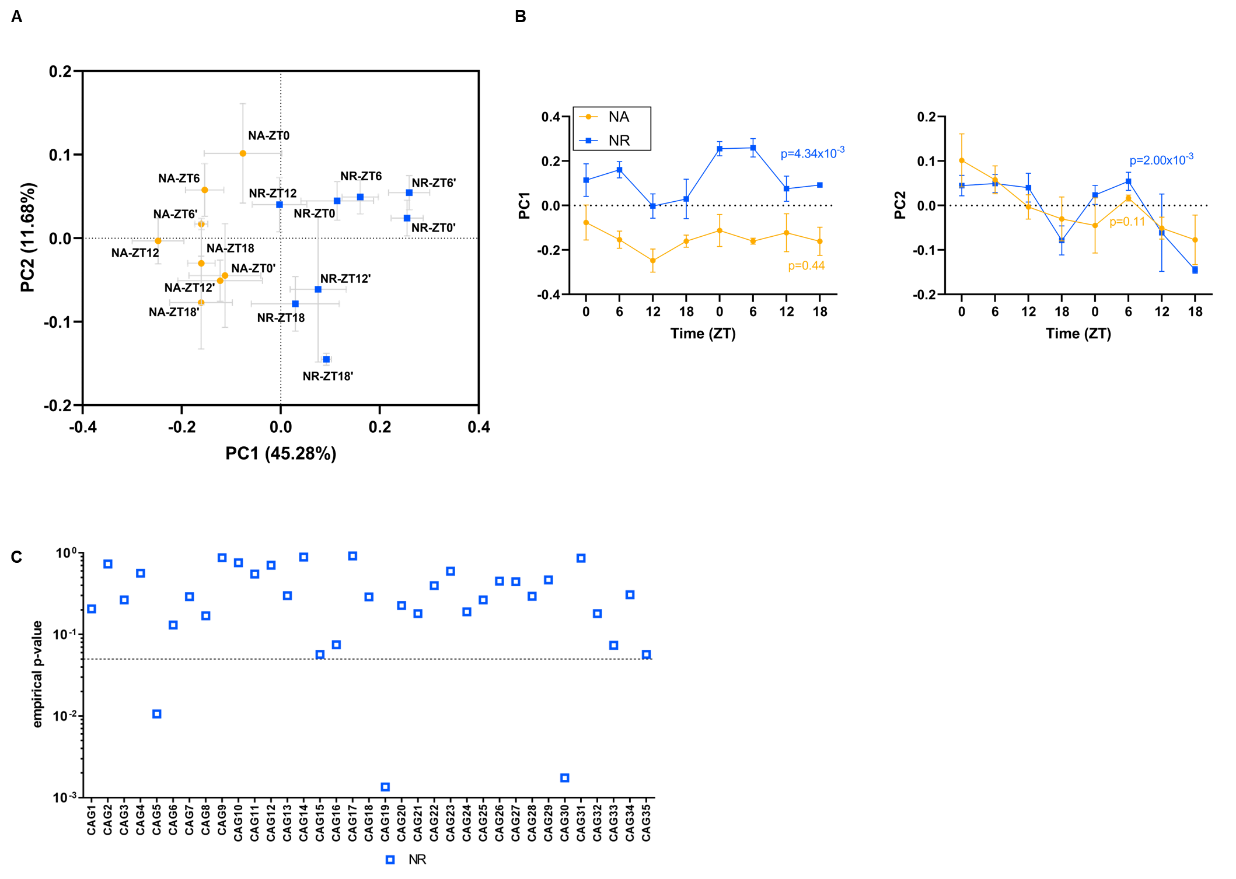

FIG S3	TRF displayed increased diurnal rhythmicity in the overall structure and composition of gut microbiota at the CAG level compared to mice with *ad libitum access to* NFD. (A) PCoA analysis was performed at the CAG level using the abundance profile of the 35 CAGs based on the Bray-Curtis dissimilarity of all the samples collected over the course of two light-dark cycles during week-11 of treatment. (B) Alteration of the gut microbiota structure along the first and second principal coordinate (PC1 and PC2) of the PCoA based on Bray-Curtis dissimilarity. Data were plotted as mean ± s.e.m.; n=3-5 fecal samples per time point. Rhythmicity was analyzed using the non-parametric empirical JTK-cycle algorithm and p-values refer to empirical-p values. (C) CAGs display diurnal oscillation in their relative abundance under NR feeding regime. Rhythmicity was examined using the non-parametric empirical JTK-cycle algorithm and p-values refer to empirical-p values. The dashed line indicates p<0.05.
