## Supplementary figures and images for "Guild-level response of the gut microbiota to isocaloric time-restricted feeding in high-fat diet-fed mice"

### Supplemental Figure 4

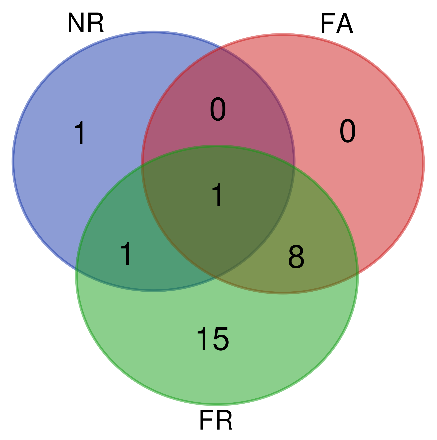
FIG S4	Number of unique and shared oscillating CAGs in each group.

### Supplemental Table 1

Table S1 Formulation of the diets used in the study

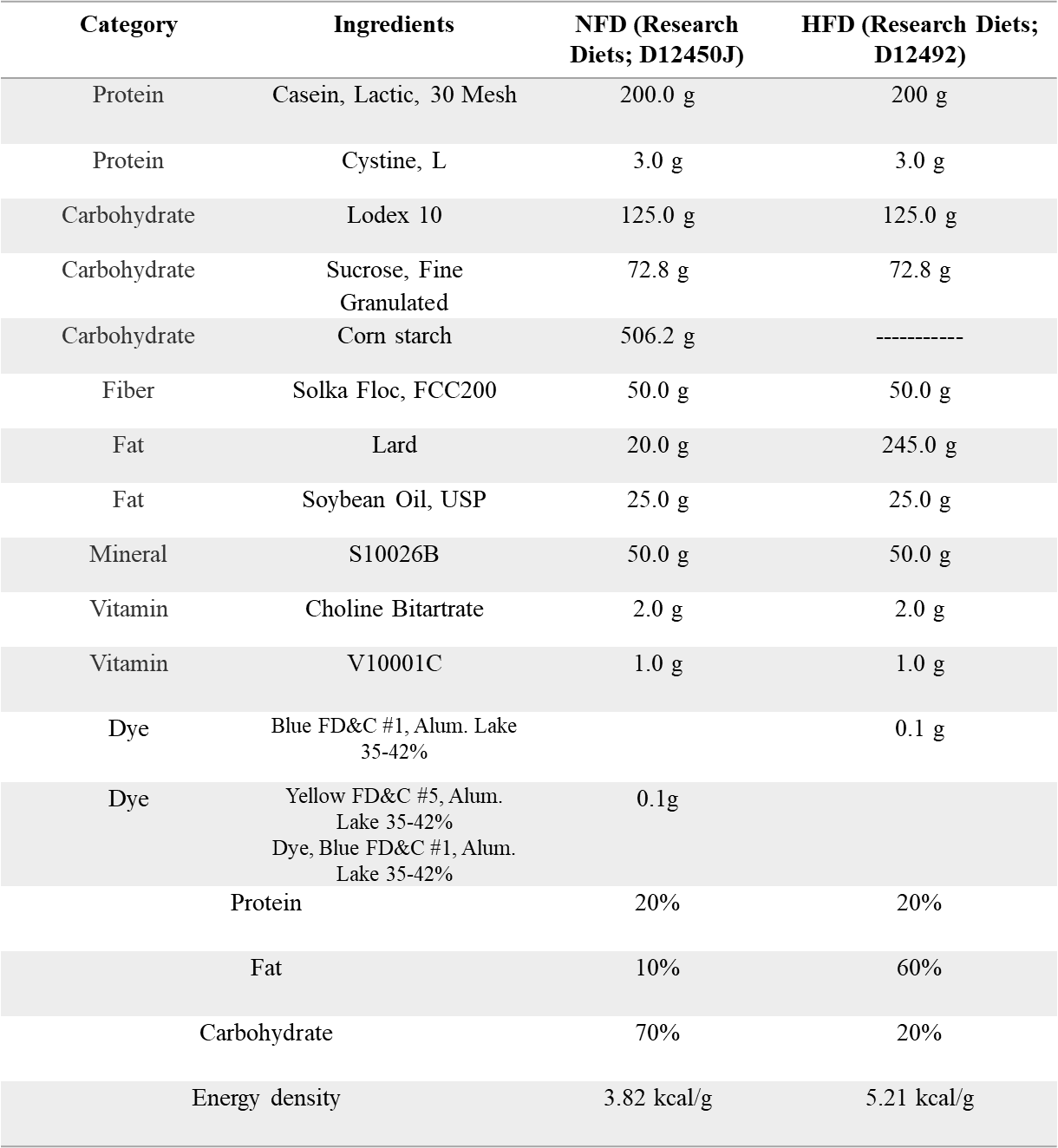

### Supplemental Table 2

Table S2	CAGs with number of ASVs with genus level annotation

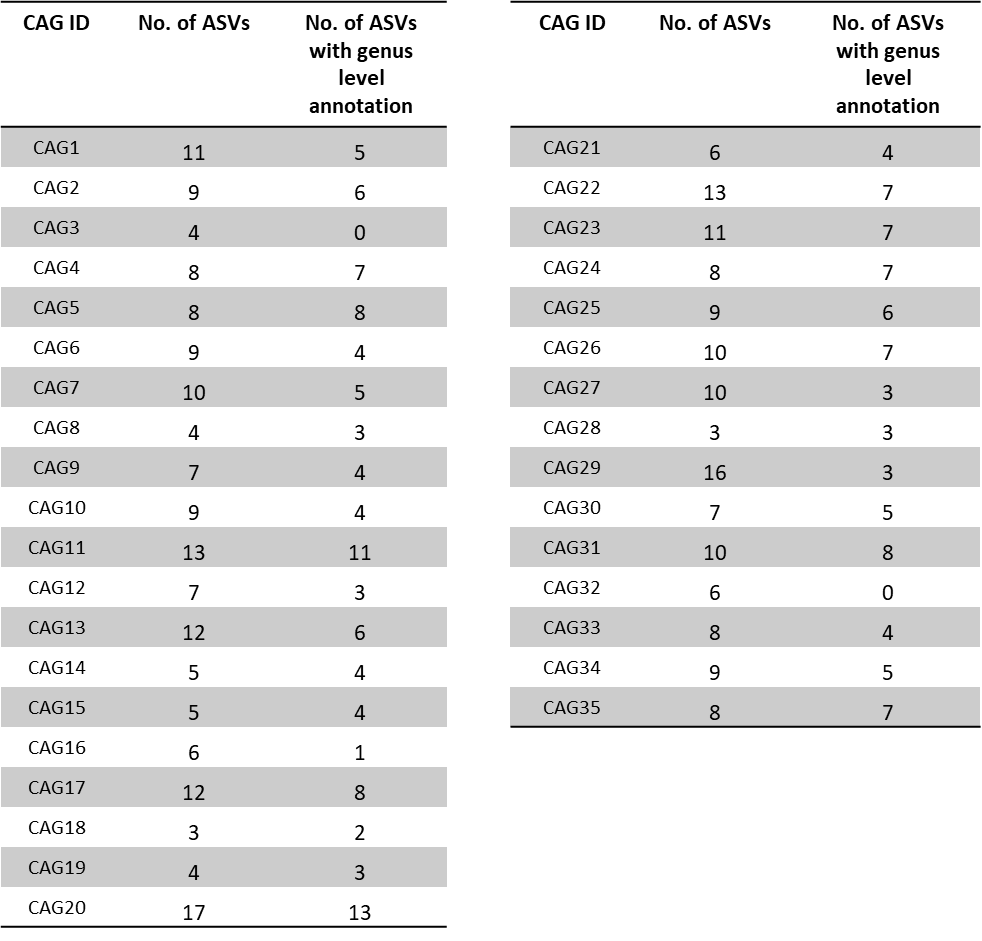

### Supplemental Table 3

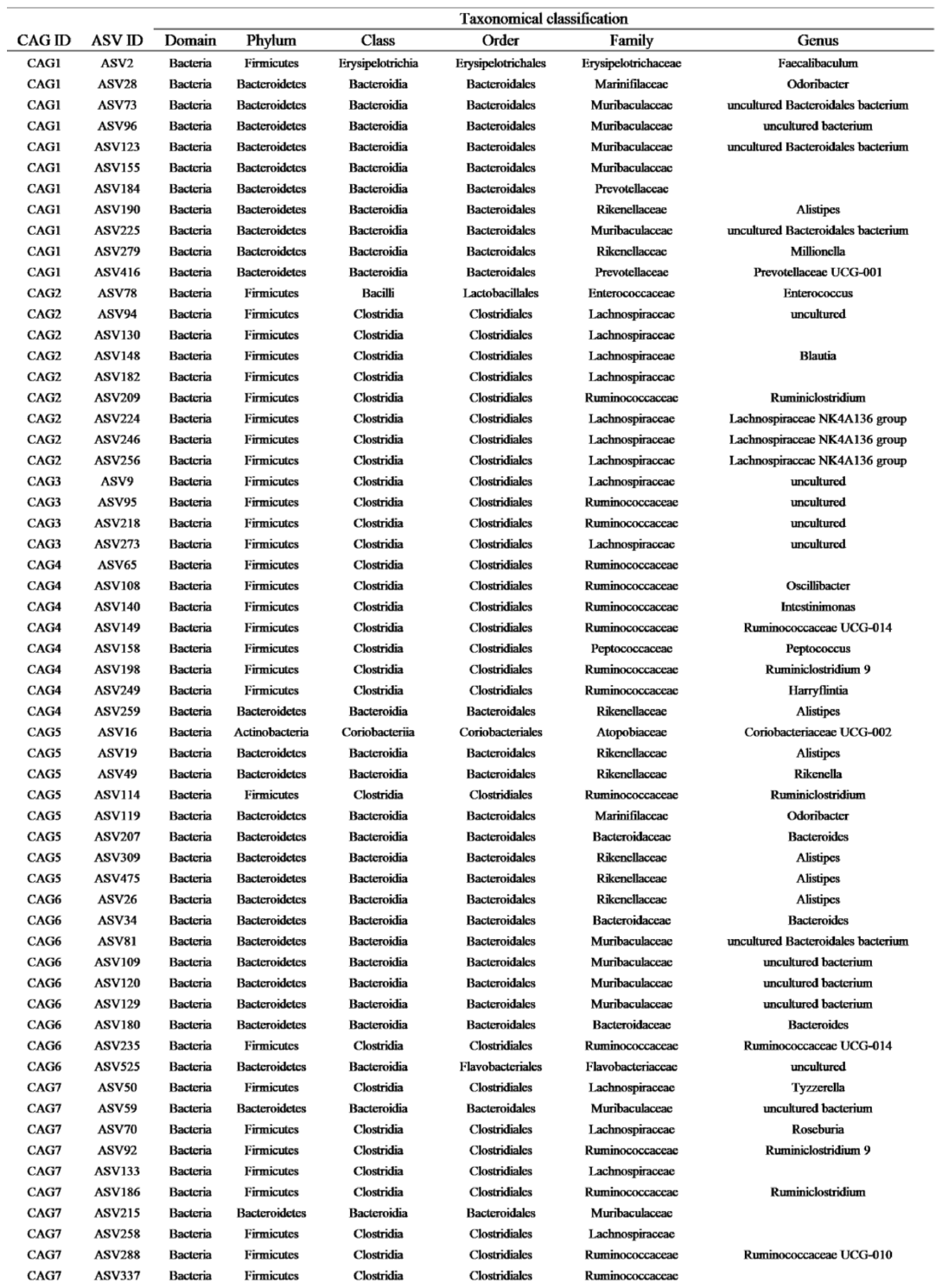
Table S3 Details of taxonomic assignments of the ASVs in the 35 CAGs

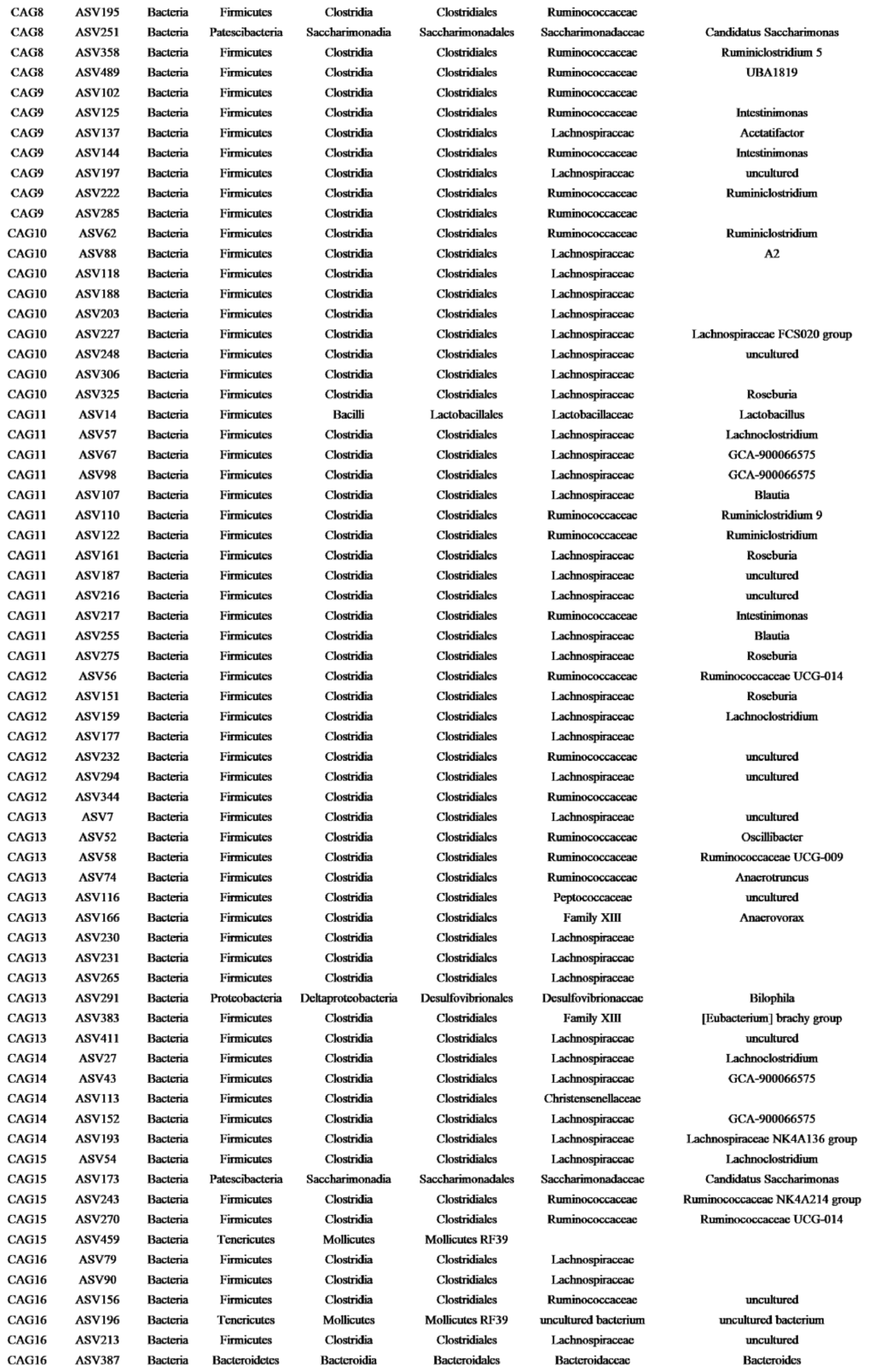

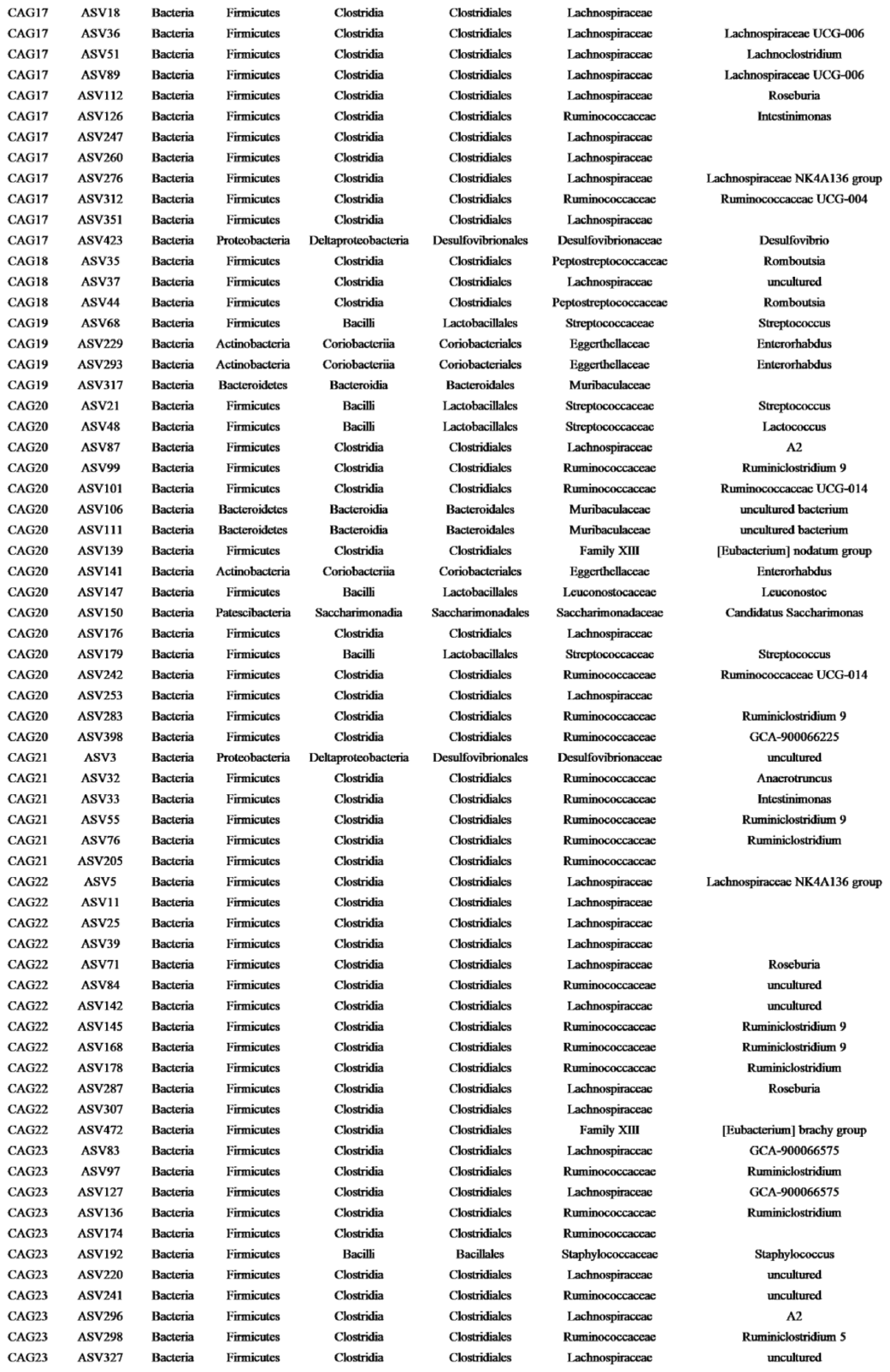

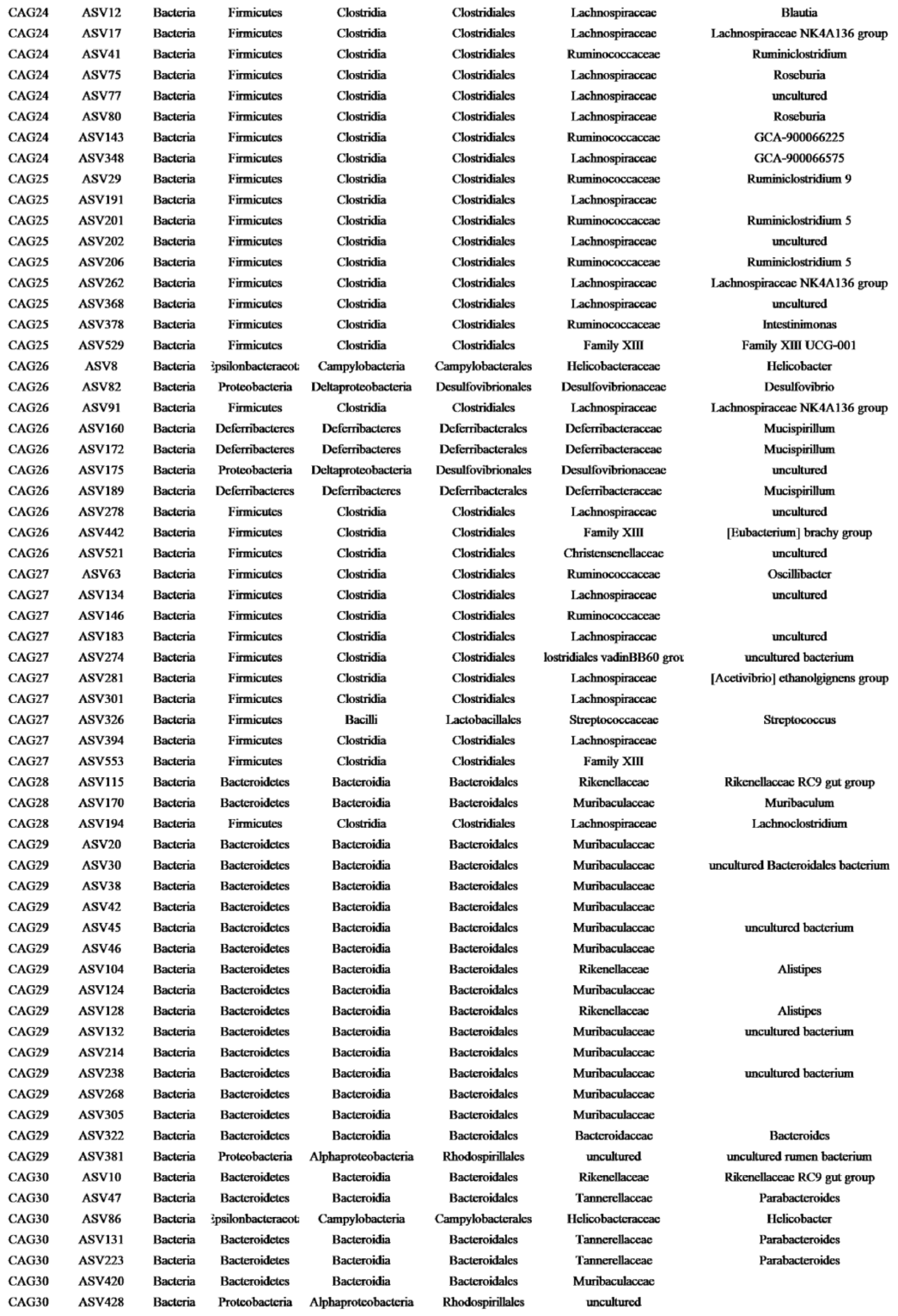

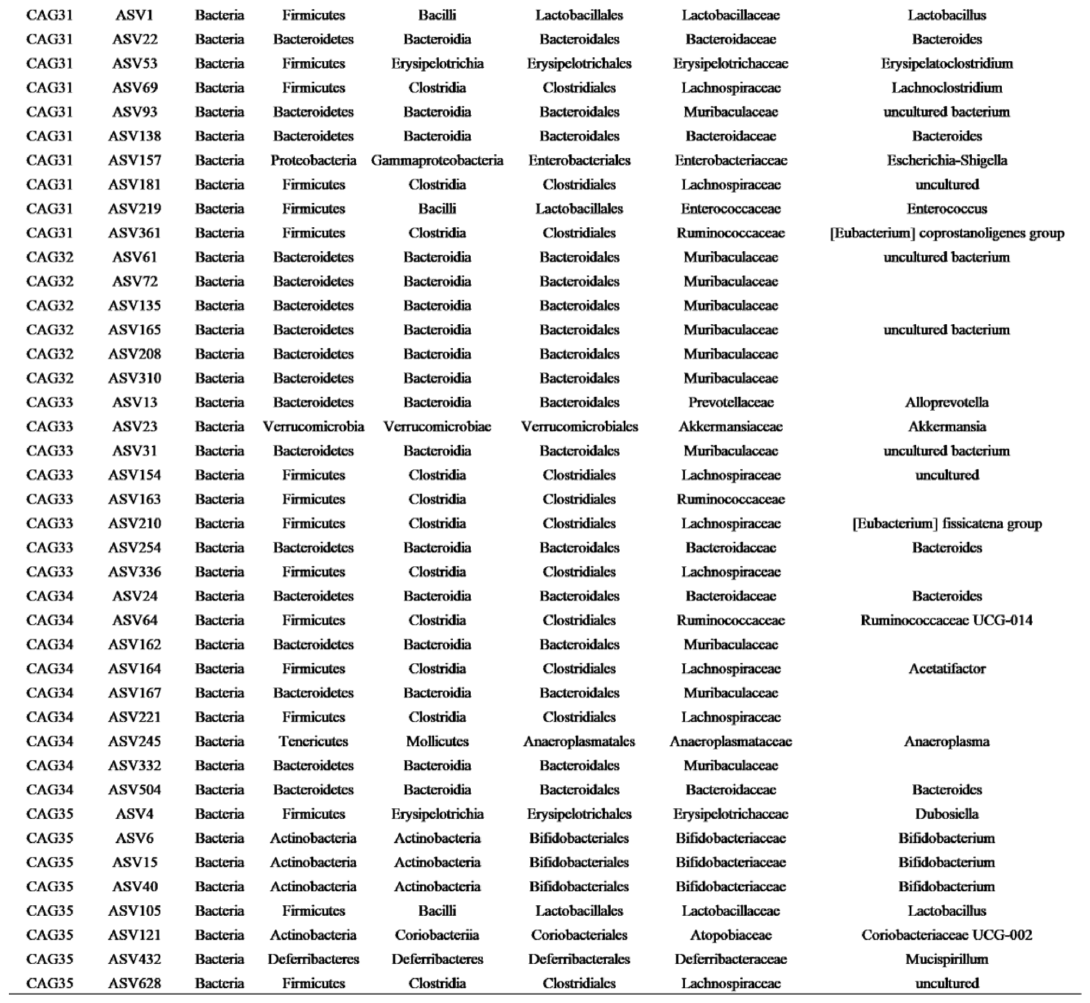
