## Supplemental Table 4 for "Guild-level response of the gut microbiota to isocaloric time-restricted feeding in high-fat diet-fed mice"

Table S4 Rhythmicity in the relative abundance of CAGs determined by empirical JTK_CYCLE in the NA and NR groups

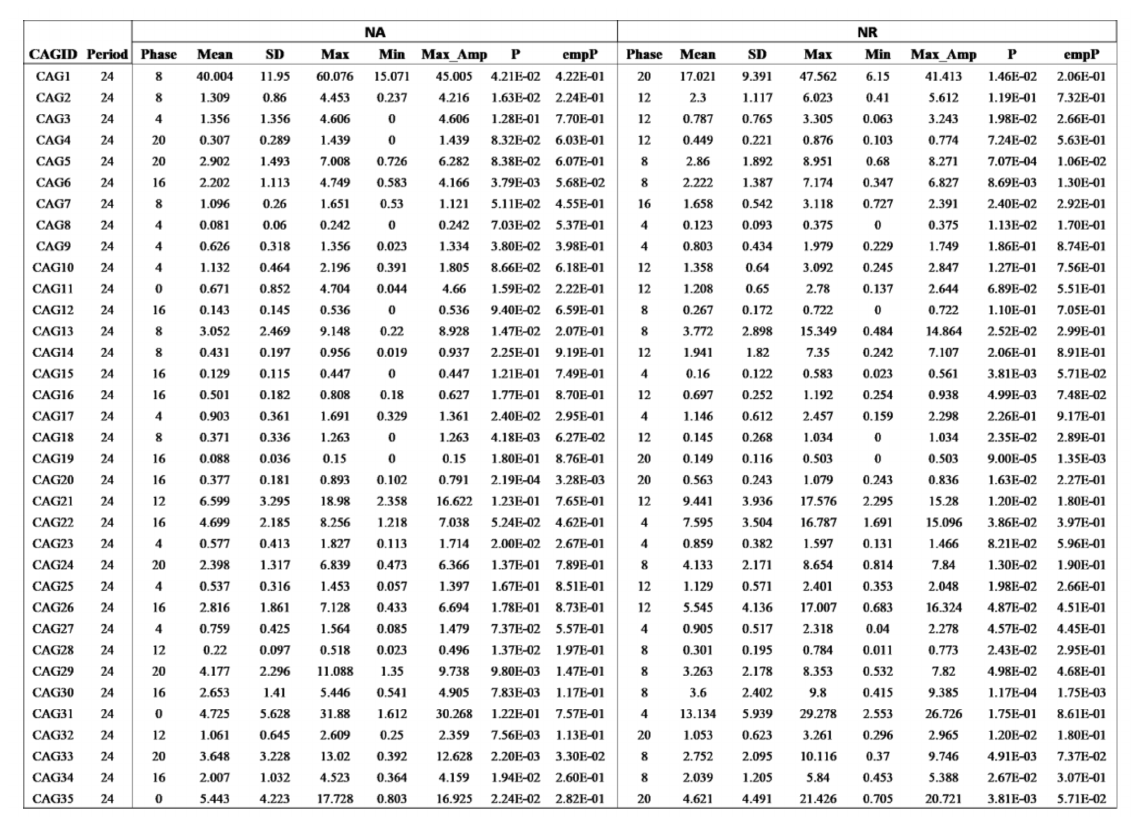
