## Supplemental Table 5 for "Guild-level response of the gut microbiota to isocaloric time-restricted feeding in high-fat diet-fed mice"

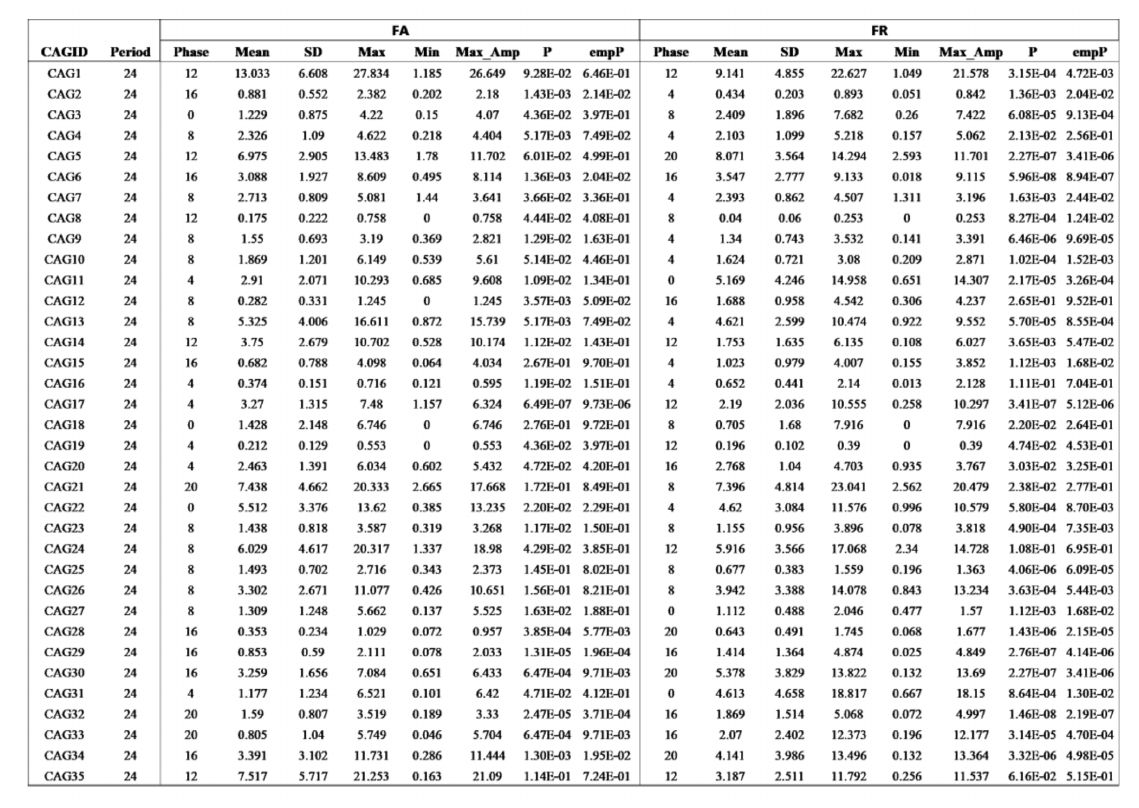
Table S5 Rhythmicity in the relative abundance of CAGs determined by empirical JTK_CYCLE in the FA and FR groups
